# Comparative framework and adaptation of ACME HS approach to single cell isolation from fresh-frozen endocrine tissues

**DOI:** 10.1101/2024.03.26.586727

**Authors:** Marina Utkina, Anastasia Shcherbakova, Ruslan Deviatiiarov, Alina Ryabova, Marina Loguinova, Valentin Trofimov, Anna Kuznetsova, Mikhail Petropavlovskiy, Rustam Salimkhanov, Denis Maksimov, Eugene Albert, Alexandra Golubeva, Walaa Asaad, Lilia Urusova, Ekaterina Bondarenko, Anastasia Lapshina, Alexandra Shutova, Dmitry Beltsevich, Oleg Gusev, Larisa Dzeranova, Galina Melnichenko, Ildar Minniakhmetov, Ivan Dedov, Natalya Mokrysheva, Sergey Popov

## Abstract

Current scRNA-seq studies of solid tissues mostly rely on enzymatic dissociation of fresh samples or the fallback on nuclei isolation from frozen or partially fixed samples. However, due to the complex tissue organization or cell fragility, it could be challenging to apply these approaches to the sensitive endocrine tissues. That is, dissociating intact cells from such problematic fresh-frozen samples routinely collected by biobanks remains challenging.

In this study, we adapted the acetic-methanol dissociation method – ACME High Salt (ACME HS) to effectively isolate intact single cells from fresh-frozen endocrine tumor samples, including adrenal gland neoplasms, thyroid carcinomas, and pituitary neuroendocrine tumors. We compared the ability of enzymatic, ACME HS, and nuclear isolation methods to preserve the integrity of major cell types and gene expression across 41 tissue samples of different origins. We demonstrated that ACME HS simultaneously dissociates and fixes cells, thus preserving morphology and a high RNA integrity number in problematic cell types. This finding renders the ACME HS dissociation method a valuable alternative in scRNA-seq protocols for challenging tissues where obtaining live cell suspension is difficult or impossible.

## Background

Recent advances in single-cell RNA sequencing (scRNA-seq) have dramatically expanded our understanding of the cellular complexity and heterogeneity of human tissues, including the endocrine glands[1],[2],[3]. However, further progress in this field struggled with the incomplete molecular characterization of the particular cell types being responsible for the functional complexity of human endocrine tissues. One of the most problematic issues in the scRNA-seq profiling of human tissues that significantly impacts the biological relevance of the ultimate data is the sample preparation step. In that, the most commonly used immediate processing of freshly isolated tissues is extremely poorly integrated in the clinical logistics of tedious surgical procedures and subsequent surgical pathology assessments. This issue is particularly important in the context of the inherent problems in obtaining the high-quality single cell suspensions from solid tissues, requiring complex disaggregation/dissociation steps while preserving the high cellular yield and viability, unbiased cellular contents, transcriptional profiles, cellular states etc. Indeed, the suboptimal procedures being employed for tissue collection, storage, and, particularly, disaggregation/cell dissociation, the procedures being associated with the excessive mechanical stress, suboptimal temperature conditions, prolonged enzymatic digestion, and the loss of the original tissue context, have been reported to dramatically distort the resulting scRNA-seq data and may even result in cell type misclassification[4],[5].

Since the late 1970s, when the first methods for the disaggregation of solid tissues were described[6],[7], a variety of protocols utilizing mechanical, enzymatic, and chemical methods of dissociation (and combinations thereof) have emerged[8],[9]. Probably the most popular approach for obtaining single cell suspensions for scRNA-seq is enzymatic digestion implying the incubation of gross tissue samples with various collagenases at 37℃ as a key step. However, a number of studies have demonstrated that the employment of these techniques is associated with the profound activation of the stress signaling pathways and increased cell death, resulting in a significant bias in the scRNA-seq profiles[5],[10]. Such undesirable effects may be largely diminished via employment of the so-called “cold” dissociation techniques with a cold active protease (6 °C)[11], however, at the cost of less efficient target cell dissociation[12].

Furthermore, the enzymatic dissociation of normal and diseased endocrine tissues may be particularly challenging due to a number of confounding structural issues, such as a high lipid content in the normal adrenal cortex and adrenocortical neoplasia, or extensive stromal/capsular fibrosis and calcifications in the well-differentiated thyroid tumors. Single-nucleus RNA sequencing (snRNA-seq) protocols being compatible with the use of the fresh-frozen tissue samples may be successfully employed to overcome these limitations[10],[13]; however, at the cost of the loss of a significant amount of the mature cytoplasmic mRNA, resulting in a lower coverage and poor representation of the rare cell types[14].

Each of the abovementioned approaches contributes to the emergence of a variety of artifacts related to the distortion of the transcriptional profiles of individual cells[12], an issue that must be explicitly addressed in the process of the ultimate data analysis[15]. For example, the immediate-early response genes (e.g., the members of the *FOS* and *JUN* gene families) are primary candidates for changing their expression during single-cell dissociation at 37°C. Artifactual changes in gene expression patterns were investigated by comparing the transcriptional profiles of cryopreserved and living cells or methanol-fixed and living cells obtained from tissue dissociation using cold-active protease and enzymatic digestion at 37 °C. This study showed that cold-active proteases dramatically reduce the number of scRNA-seq artifacts in the mouse kidney[16]. However, addressing of these issues remains rather fragmentary and limited to certain tissue types and protocols, so numerous artifacts still need to be confidently addressed.

Many of the limitations of the currently employed techniques may be potentially overcome via simultaneous tissue dissociation and cell fixation, the procedure being capable of maintaining a high RNA Integrity Number (RIN) while minimizing the sample preparation-related distortions in the transcriptional profiles. Recently, García-Castro et al.[17] introduced such a protocol whose prototype may be tracked back to the end of the 19th century[18],[19], when Schneider reported the so-called “maceration” technique. In its contemporary variant, named ACME (ACetic (acid)-MEthanol), this technique reportedly produces the high-quality suspensions of fixed single cells from planarians, *D. melanogaster*, *D. rerio*, and *M. musculus* tissues, the suspensions maintaining the high-integrity RNA that may be further successfully cryopreserved using DMSO[17].

Here we extensively optimized and successfully implemented the unique ACME High Salt (ACME HS, see *Methods*) protocol for the single-cell transcriptomic analysis of human neoplastic endocrine tissues represented by tumors arising from the adrenal medulla, adrenal cortex, pituitary gland, and thyroid follicular cells. We also compared our modified ACME HS and enzymatic dissociation methods for scRNA and nuclei isolation for snRNA profiling in terms of the number of the cells/nuclei recovered, RNA integrity, aligning of the resulting scRNA/snRNA data with the reference organ-specific profiles, and representation of the specific cell types.

We clearly demonstrated that scRNA profiling of single cell suspensions obtained using both methods significantly outperformed snRNA profiling in terms of marker genes expression analysis and tumorgenesis while demonstrating in-between comparable performances in virtually all implemented analyses. Additionally, the modified ACME HS protocol allows successful cryopreservation of dissociated/fixed cells without sacrificing the mRNA yield and integrity. To our knowledge, this is the first report on successful implementation of the ACME HS technique in primary human tissues, and we believe that this protocol should significantly promote the scRNA studies in humans that are to be explicitly compliant with the real-life infrastructure and logistics of the surgical care centers.

## Results

### ACME HS-based dissociation of endocrine tumor samples produces fixed cells with high RNA integrity and preserved morphology

Using the adrenocortical tumor sample, we demonstrated the morphology and found that the RNA integrity of ACME HS-dissociated adrenocortical cells was well-preserved. For ACME HS dissociation, a fresh adrenocortical tumor was previously cryopreserved in a biobank (**Fig. 1d**, 1 day) and the cell suspension obtained the next day was divided into 7 aliquots (6 aliquots for the RNA integrity number (RIN) calculation and one aliquot for microscopy). Enzyme-dissociated cells were obtained from the same fresh adrenocortical tumor (**Fig. 1d**, 0 days) and were also divided into 7 aliquots.

**Fig. 1:**
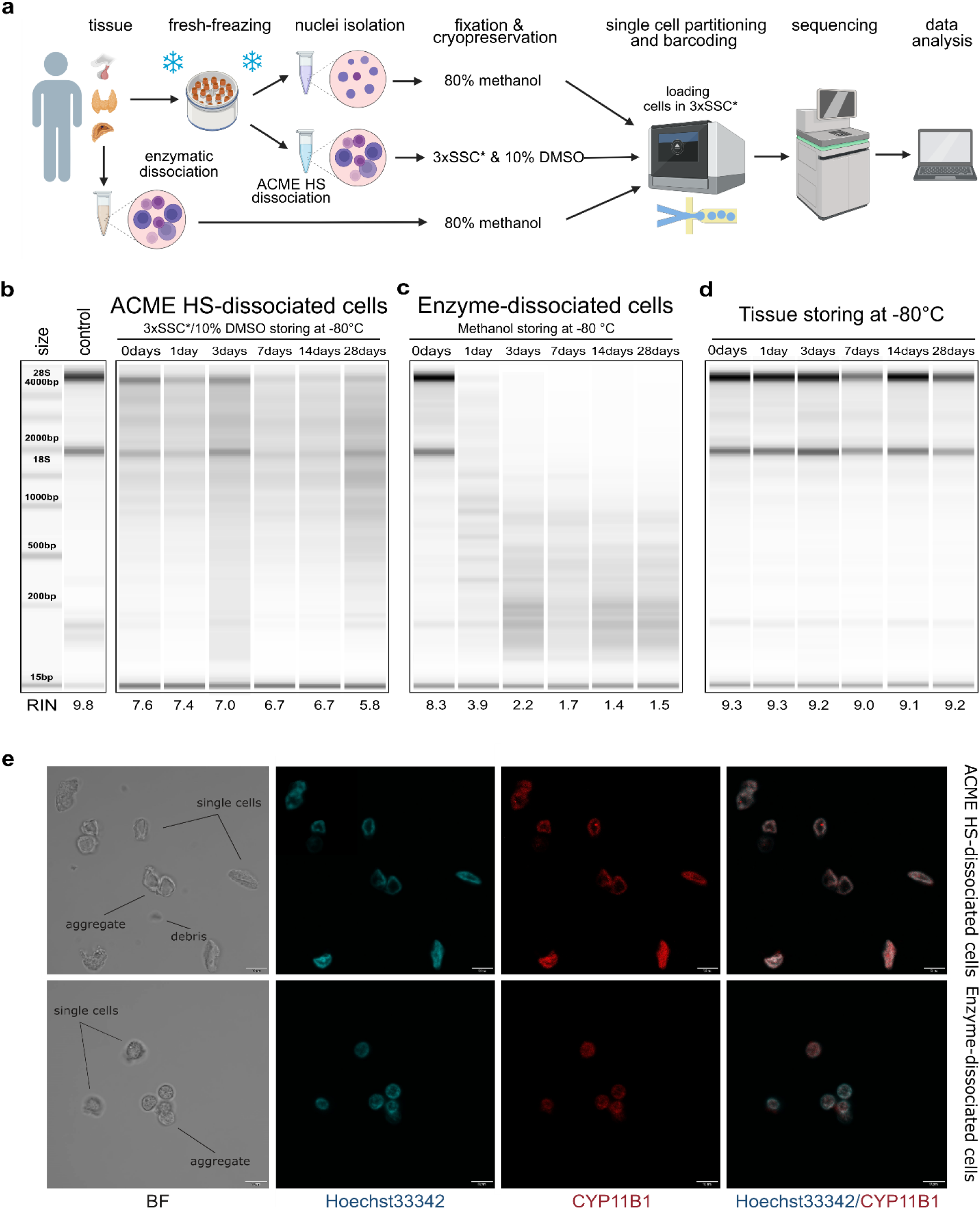
Comparison of RNA integrity, morphology, and storage of ACME HS and enzyme-dissociated adrenocortical cells. **a.** Schematic representation of a workflow for single-cell or single-nuclei processing and analysis of fresh and fresh-frozen tissues (created with *BioRender.com*). **b.** Gel image of isolated total RNA from cryopreserved ACME HS-dissociated adrenocortical cells after 1,3,7,14, and 28 days of freezing at -80°C, and of freshly isolated adrenocortical cells kept at +4°C (0 days). **c.** Gel image of isolated total RNA from adrenocortical cells obtained by the enzymatic dissociation and fixed in 80% methanol after 1, 3, 7, 14, and 28 days of freezing at -80°C, and of freshly isolated cells kept at +4°C (0 days). **d.** RNA integrity of fresh-frozen adrenocortical tumor, from which ACME HS-dissociated cells (Fig. 1b) were obtained. **e.** Bright field (BF) and confocal fluorescence microscopy images of freshly isolated ACME HS and enzyme-dissociated adrenocortical cells stained with Hoechst 33342 (blue) and anti-CYP11B1 antibody (red), showing single cells, aggregates, and debris.

Single cells were isolated from tissue samples using ACME HS and enzymatic dissociation methods, freshly or followed by cryopreservation in 3xSSC*10% DMSO and methanol cell fixation, respectively (**Fig. 1a, Additional file 2: Table S1**). The key adaptations of the ACME HS method were the supplement of the solution composition with 0.1M NAC and the introduction of additional washing steps in cold high salt 3xSSC* buffer. ACME dissociation was conducted in ∼ 1 hour on a rotator at room temperature, with periodic pipetting of the solution. Afterward, we removed the ACME solution and washed the pellet with a two-step washing in cold 3xSSC* (see *Methods* for details).

Total RNA extracted from freshly prepared cell suspensions (0 days) showed that the RINs for cells obtained through the ACME HS dissociation method and enzymatic digestion, were approximately the same, at 7.6 and 8.3, respectively. The two major ribosomal RNA subunits, 18S and 28S, were determined (**Fig.1 b, с**). Similar RINs scores of RNA were estimated for cells obtained from adrenal medullary tumor, thyroid carcinoma and pituitary neuroendocrine tumor (PitNET) samples (**Additional file 1: Figure S1b**). The obtained RIN scores were compared with the RIN score (9,8) of undissociated adrenocortical tumor (control) (**Fig. 1b).**

Next, we visualized freshly prepared dissociated adrenocortical cells (0 days) by bright field and confocal microscopy. We found that the cells preserved their morphology and exhibited minimal aggregates and debris (**Fig. 1e**). Microscopy was also performed for thyroid cells and one replicate of adrenocortical cells (**Additional file 1: Figure S2b**).

To identify adrenocortical cells, we stained the fixed cells with Hoechst 33342 and an anti-CYP11B1 (11β-hydroxylase) antibody conjugated with Alexa Fluor 594 (**Fig. 1e, Additional file 1: Figure S2b**). CYP11B1 is localized in the inner mitochondrial membrane and is normally expressed in the zona fasciculata of the human adrenal cortex[20]. We observed intense immunofluorescence of CYP11B1 (red) in ACME HS-dissociated adrenocortical cells, unlike in cells isolated using enzymatic digestion. This difference could be attributed to the prolonged permeabilization of cell membranes with methanol during the ACME HS protocol. The voids formed in the nuclei of adrenocortical stained cells are likely associated with their functional ability to efflux the DNA binding dye Hoechst 33342, resulting in the so-called side population (SP)[21]. Thyroid follicular cells were also visualized by staining fixed cells with Hoechst 33342 and an anti-TSHR (TSH receptor) antibody conjugated with Alexa Fluor 594 (**Additional file 1: Figure S2b**).

### ACME HS-di2ssociated cells can be cryopreserved and stored

A key disadvantage of commonly used enzymatic digestion methods is that freshly isolated tissues are extremely poorly integrated in the clinical logistics of surgical procedures. Importantly, enzyme-dissociated cells can only be cryopreserved or fixed after dissociation, typically with DMSO or methanol. Under such protocols, however, cells remain outside their extracellular space, for an extended period, and fixation in methanol leads to further structural changes and artifact introduction. Additionally, some economic constraints are associated with the need to fully load the Chromium Next GEM chip with eight samples for cell capture and barcoding.

The method we used made it possible to freeze the fresh tissue sample for subsequent ACME HS dissociation. Since the ACME HS method simultaneously carries out tissue dissociation and cell fixation, the resulting suspensions can be cryopreserved for subsequent analysis[17]. To confirm that the ability of cryopreserved of ACME HS-dissociated cells in 3xSSC* and 10% DMSO allows to maintain RNA integrity, we compared ACME HS-dissociated cells after various freeze periods.

We sequentially extracted total RNA from 6 aliquots of ACME HS and 6 aliquots of enzyme-dissociated cells obtained from the same adrenocortical tumor (**Fig. 1d**) at different time intervals, starting from the moment of the freshly prepared single-cell suspension (0 days) and after 1, 3, 7, 14, and 28 days of cryopreservation and methanol fixation (**Fig. 1b, c**). The obtained RIN scores were compared with the RIN scores of the control (**Fig. 1b**).

We observed that the RNA integrity of ACME HS-dissociated cells was maintained during the cryopreservation in 3xSSC* and 10% DMSO over the specified time intervals (1, 3, 7, 14, and 28 days). RIN was measured in each case giving a score ∼ 6.7 (**Fig. 1b**). As a result, the cryopreservation of ACME HS-dissociated cells obtained from adrenocortical tumor as well as adrenal medullary tumor, thyroid carcinoma and PitNET in 3xSSC*/10% DMSO maintains RNA integrity for subsequent scRNA-seq analysis (**Fig. 1b, Additional file 1: Figure S1**). Furthermore, the RNA integrity of cryopreserved ACME HS-dissociated adrenocortical cells after six months of storage was 5.9 (**Additional file 1: Figure S2a**). The RNA integrity of the dissociated cells obtained from PitNET was estimated for only two time intervals (0 days and 1 day) due to the small size of the tissue sample.

However, the RIN of enzyme-dissociated adrenocortical cells fixed in 80% methanol decreased over the specified time intervals (1, 3, 7, 14, and 28 days). The observed degradation of ribosomal RNA was reflected in the reduction of signal intensity or its complete absence for both ribosomal peaks (18S and 28S) (**Fig. 1c, Additional file 1: Figure S1b**).

The RNA integrity of a fresh-frozen adrenocortical tumor samples after 1, 3, 7, 14, and 28 days of freezing at -80°C ranged from 9.3 to 9.0. No discernible patterns in the change of RIN score over time were identified (**Fig 1d**). The obtained RIN scores indicated the suitability of the material for subsequent dissociation using the ACME HS method.

### Flow cytometry reveals heterogeneity of ACME HS and enzyme**-**dissociated endocrine samples

Flow cytometry was used to assess the quality of ACME HS and enzyme-dissociated samples. We analysed various tissues, including adrenocortical tumors, adrenal medullary tumors, thyroid carcinomas, and PitNETs. We compared the amount of debris, aggregates, and single cells obtained by the two protocols of tissue dissociation – ACME HS and enzymatic. (**Fig. 2, Additional file 1: Figure S3, replicate 1-3**). It appeared more straightforward to calculate the amount of cellular debris and the number of singlets and aggregates at the FSC-area/FSC-height dot plot (**Fig. 2, a1**) than at the FSC/SSC dot plot (**Fig. 2, a2**), as the boundary between debris and singlets, and between singlets and their aggregates, is usually poor. We gated out the area of debris (green) as small events with the highest ratio of FSC-height to FSC-area signal. Singlets (red) were selected based on their well-correlated height versus area signal, while aggregates of cells (black) had an increased area signal compared to the height signal. (**Fig. 2, a1; Additional file 1: Figure S2, a1-h1**).

**Fig. 2:**
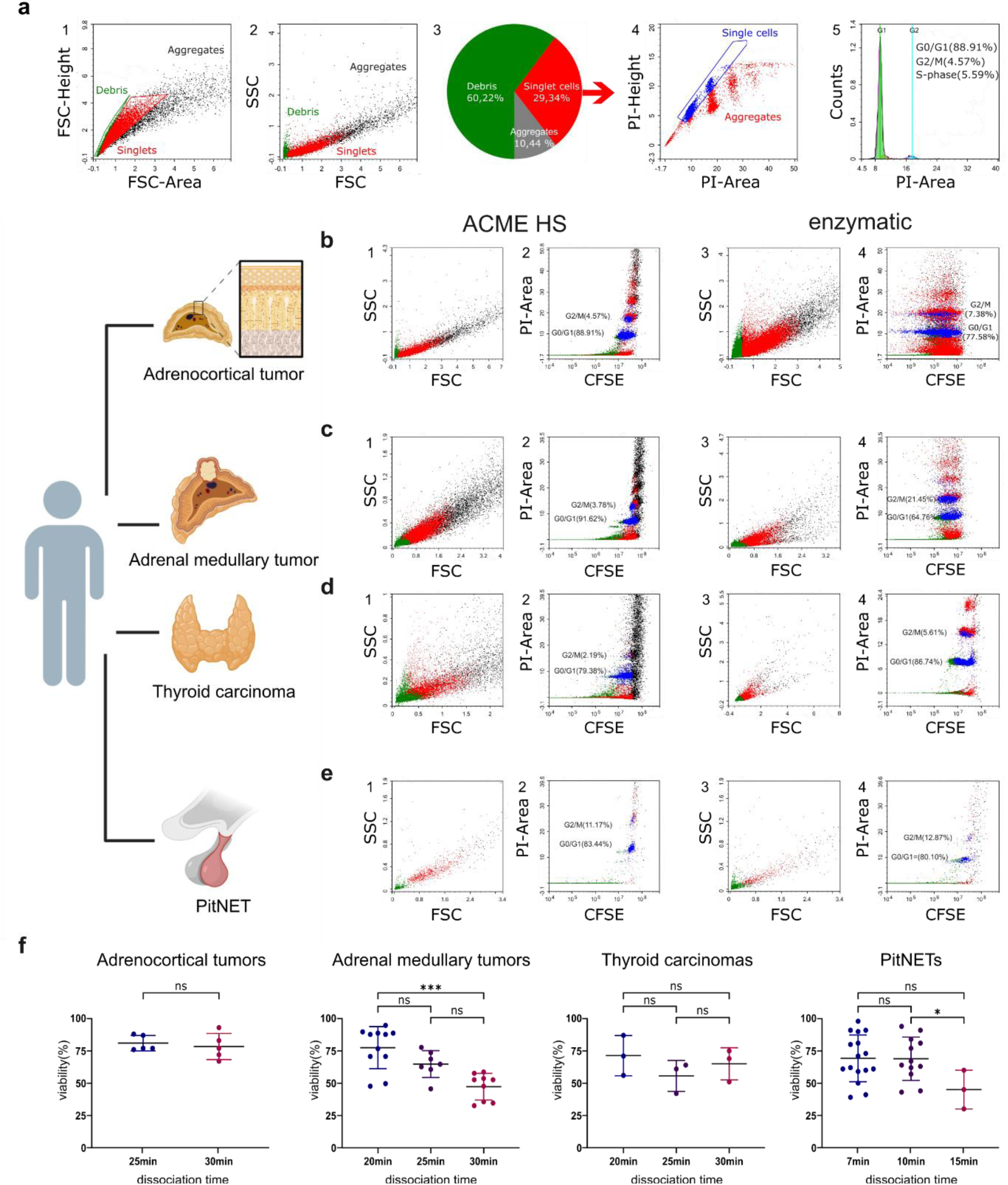
Representative flow cytometry data of the samples prepared by ACME HS and enzymatic dissociation methods. **a.** Flow cytometry data for the adrenocortical tumor sample (replicate 1). **a1.** FSC-height/FSC-area dot plot was used to calculate cellular debris, single cells, and cellular aggregates (green events - for debris, red events - for singlets, black events – for aggregates). **a2.** FSC/SSC dot plot demonstrating the distribution of cells, their aggregates, and cellular debris based on their light-scattering properties. **a3.** Pie diagram of the debris, singlets and aggregates distribution in the sample. **a4.** PI-height/PI-area dot plot of singlets used for additional gating of single cells (shown in blue) among nucleated cells and their aggregates (shown in red). **a5.** DNA histogram from single events showing the cell cycle distribution for all cells in the sample, with percentages of the cell cycle phases (G0/G1, S, G2/M) inserted. **b.** Flow cytometry data for the different samples (replicate 1). **b1-b4** – for the adrenocortical tumor; **c1-c4** – for the adrenal medullary tumor; **d1-d4** – for the thyroid carcinoma; **e1-e4** – for PitNET (green events - for debris, red events - for singlets, black events – for aggregates). Index 1 stands for FSC/SSC dot plots for the samples obtained by the ACME HS protocol. Index 2 stands for CFSE/PI dot plots for the samples obtained by the ACME HS protocol. Index 3 stands for FSC/SSC dot plots for the samples obtained by enzymatic dissociation. Index 4 stands for CFSE/PI dot plots for the samples obtained by enzymatic dissociation. **f.** Cell viability (%) data of adrenocortical tumors, adrenal medullary tumors, thyroid carcinomas, and PitNETs cells at different dissociation times. The values are plotted for each experiment, and the mean ± SEM is indicated. Statistical significance was estimated by t-test: *** (*p* < 0.001),* (0.01 < *p* < 0.05), ns - not significant – *p* > 0.05. (Created with BioRender.com).

Further, we used a strategy to evaluate the data in other dot plots to correct our subsequent gating for the best separation and calculation of the amount of cellular debris and the number of singlets and aggregates. First, we applied color gating and visualized debris, singlets and aggregates in the FSC/SSC dot plots for all tissues (**Fig 2, a2, b1-e1, b3-e3; Additional file 1: Figure S3,a2-h2**).

We made sure that debris was located in the lower left area of the dot plot, and the aggregates formed clusters in the upper right area of the FSC/SSC dot plot. Next, we stained the samples with a DNA-binding dye – propidium iodide (PI), to better discriminate the nucleus-contained cells from the nuclear-free debris. We studied the PI-height/PI-area dot plots and checked the positions of debris, singlets, and aggregates among all ungated events including debris and aggregates (**Additional file 1: Figure S3, a3-h3**). Then, we gated the single cells (blue) among the nucleated cells and their aggregates (red) (**Fig. 2, a4; Additional file 1: Figure S3, a4-h4**) and analysed DNA histograms from single cells (**Fig. 2, a5; Additional file 1: Figure S3, a5-h5**).

Although our dissociation protocols differed from those that were specifically elaborated for cell cycle analysis and frequently used[22], in most cases, we could resolve various phases of the cell cycle (G0/G1, S, G2/M) by mean fluorescence intensity (MFI) per cell in DNA histograms: DNA content in G2/M phase was as expected, two times more than in G0/G1 as shown in **Additional file 1: Figure S3, a5-h5**.

To discriminate the nature of the debris, we stained the samples with 5,6-carboxyfluorescein diacetate succinimidyl ester (CFSE). This dye is frequently used in flow cytometric protocols for live cells labeling due to its ability to bind to intracellular molecules, primarily to amine groups. In addition to its role in viable cell staining, CFSE can trace dying cells in composite samples[23]. As shown in **Fig. 2, b2-e2** and **b4-e4**, most green events matching debris turned out to be CFSE-positive and PI-negative, which suggested that the debris was generally nuclear-free. By comparing the frequency of debris and aggregates (**Additional file 1: Figure S3, a1-h1**) and analyzing dot plots, we suggest that both ACME HS and enzymatic methods induced a relatively similar number of aggregates and debris. Despite the large amount of debris and aggregates debris and aggregates, which was expected, we observed a sufficient number of single cells in our samples obtained by the ACME HS and enzymatic dissociation protocols (**Fig. 2, a2; Fig. 2, b1-e1, b3-e3; Additional file 1: Figure S3, a2-h2**).

### Effect of enzyme dissociation protocols on the cell viability of endocrine tumor samples

We investigated the impact of tissue dissociation time and enzyme type on the viability of cells isolated from adrenocortical and adrenal medullary tumors, thyroid carcinoma, and PitNET samples. Various enzymes, including collagenase I, collagenase IV, MTDK, and NTDK, were used for tissue digestion (see *Methods*). We also tested different dissociation times. In this assay, we included samples corresponding to the tissues under study and were not identified as blood (**Additional file 2: Table S1**).

We observed no significant differences in the impact of enzyme type on cell viability (**Additional file 1: Figure S2c**). To assess the impact of dissociation time on the samples, cell viability was compared after 20, 25, and 30 minutes of enzyme dissociation for adrenal medullary tumor (n=26) and thyroid carcinoma (n=9) samples; after 25 and 30 minutes for adrenocortical tumors (n=10); and after 7, 10, and 15 minutes for PitNET (n=20) samples, using the trypan blue staining assay (**Fig. 2f**).

A wide range of cell viability was noted for adrenal medullary tumor samples at 20 and 30 minutes of dissociation, showing significant differences (p=0.0001, t-test) with means of 77.8 and 47.7, respectively (**Fig. 2f**). Thus, the highest cell viability was achieved with a 20-minute dissociation for adrenal medullary tumor samples. Our findings suggest a compromised cell viability (approximately 50%) for the 30-minute dissociation protocol. Finally, significant differences (p=0.0449, t-test) were identified for the PitNET samples at 10 and 15 minutes of dissociation time, with means of 69.2 and 45, respectively. This indicates that the dissociation time should not exceed 10 minutes. It is important to note that the transnasal removal of the pituitary gland often leads to tumor fragmentation. Additionally, irreversible changes during dissociation can occur abruptly with even minor variations in incubation time or enzyme concentration.

No correlation was found between enzymatic dissociation time and the viability of adrenocortical and thyroid follicular cells (**Fig. 2f**). It should be noted that a larger volume of MTDK (see *Methods)* was used for the enzymatic dissociation of thyroid carcinomas than for the dissociation of other tissues. Areas of fibrosis, amyloidosis, and calcinosis may occur in tumor tissues, making dissociation difficult. In particular, thyroid carcinoma contains a large amount of collagen and a colloid, which is a homogeneous gelatinous substance.

Although we sought to determine the influence of tissue dissociation time and enzyme type on cell viability, external factors cannot be excluded. Specifically, apoptosis and cell necrosis, coagulation changes, and secondary tissue changes such as fibrosis, cholesterol crystal deposition, cystic transformation, and varying-duration hemorrhages may influence dissociation (**Additional file 1: Figure S4**). Thus, due to the individual and poorly predictable properties of each sample, the dissociation time can vary between 3 and 15 minutes.

### ACME HS demonstrates consistency with enzymatic method and keep advantages over nuclei isolation protocol

To further perform comparative analyses, we selected 41 human endocrine tumor samples (**Additional file 2: Table S1**). We obtained 107.875 cells and nuclei isolated from adrenocortical tumors (n=12), 94.807 cells and nuclei from adrenal medullary tumors (n=15), 41.418 cells and nuclei from PitNETs (n=9), and 60.365 cells from thyroid carcinomas (n=5).

First, we examined the summary statistics for the generated single cell gene libraries (**Fig. 3a, Additional file 1: Figure S5**). We found that the quality of the ACME HS dataset was almost identical to the enzymatic and nuclei datasets, obtained from ACME HS-dissociated and enzyme-dissociated whole cells, and isolated nuclei, respectively. Some differences in ribosomal and mitochondrial gene expression levels between nuclei and whole cells obtained by ACME HS and enzymatic dissociation methods were confirmed directly, as well as the exon/intron alignment ratio (**Fig. 3a**). At the same time, there were no significant differences in quality parameters, namely the total number of cells, reads in cells, ambient RNA and doublets fraction for all samples (**Fig. 3a**). We observed the patterns mentioned earlier in the four tissues with different dissociation methods (**Additional file 1: Figure S5**).

**Fig. 3:**
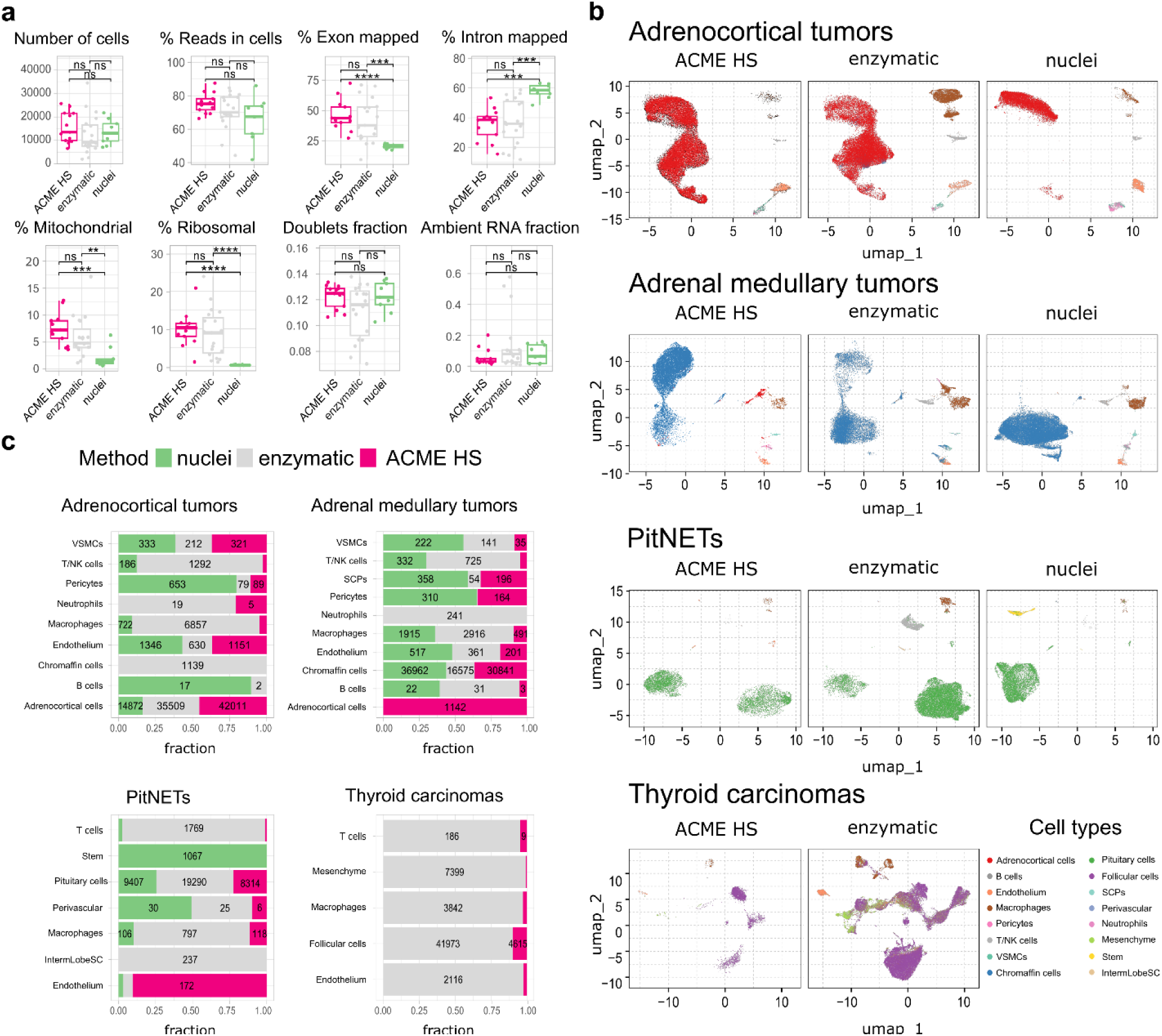
ACME HS demonstrates consistency with the enzymatic protocol and keeps advantages over the nuclei isolation. **a.** Standard single-cell sample features (number of cells; read the cells%; exon/intron mapped; mitochondrial, ribosomal, and ambient RNA; doublets) of ACME HS, enzyme and nuclei samples. Statistical differences estimated by the Wilcoxon rank-sum test: **** (0.0001 < *p* < 0.001), *** (*p* < 0.001), ns - not significant – *p* > 0.05. **b.** Major cell type compositions among preparation methods, namely adrenocortical, chromaffin, pituitary, and thyroid follicular cells. **c.** Fractions of defined cells identified by different dissociation methods (ACME HS, enzyme, nuclei) for each tissue type – adrenocortical tumor, adrenal medullary tumor, thyroid carcinoma, and PitNET. The diagram does not indicate the number of cells representing less than 5% of the total number.

To compare the representation of distinct cell types and states between the enzyme, nuclei, and ACME HS datasets, we used Seurat[24] to generate integrated embeddings and annotations for each tissue type. The major cell types were successfully defined and integrated via all three methods. In all four types of tissues, the ACME HS data retained tissue-specific cells, namely adrenocortical, chromaffin, pituitary, thyroid follicular cells, and other nonspecific cells (**Fig. 3b, c**). The heterogeneity of the major cell populations (adrenocortical, chromaffin, thyroid follicular, and pituitary cells) was estimated by further clustering of the integrated cells (**Additional file 1: Figure S6**). Minor subclusters (<100 cells) were excluded from the analysis. We found that most subclusters (A-2, A-3, A-5, A-4, A-7, A-18, and A-19) were lost for the adrenocortical samples in the nuclei datasets, unlike in the ACME HS and enzyme datasets.

Loss of numerous subclusters was also observed in the nuclei datasets of the chromaffin and pituitary samples (C-2, C-4, C-8, C-20, C-23, and C-25 and P-1, P-2, P-3, P-5, P-10, P-15, P-16, and P-17, respectively). However, some subclusters were more enriched in the nuclei datasets for chromaffin samples (С-0, С-1, С-5, С-6, С-14, and С-19) (**Additional file 1: Figure S6**).

While some cell subpopulation variability is expected due to the individual tissue conditions, the most apparent difference is determined for nuclei-based samples. The main clusters of adrenocortical subpopulations – A-3, A-4, A-5, and chromaffin cells – C-2 C-3 C-4 (**Additional file 1: Figure S6a**) enriched in oxidative phosphorylation, electron transport chain, ribosomal, mitochondrial, and mRNA processing genes are missing from nuclear datasets as opposed to ACME HS and enzymatic samples **(Additional file 1: Figure S7a, b**). Because of the low cell numbers obtained from thyroid carcinoma samples with ACME HS-dissociation, comparisons of major thyroid follicular cell population clusters obtained with ACME HS and enzymatic dissociation methods were not possible for these samples.

### ACME HS-dissociated cells maintain the expression of key tissue-specific genes

We next examined whether the extent to which ACME HS dissociation was able to preserve the expression of tissue-specific marker genes in all four tissue samples. In total, 107.875 cells were obtained from adrenocortical tumor samples (n=12) and integrated. Adrenocortical cells accounted for 85.6% of the annotated cell types for (**Fig. 3c**). These cells expressed key literature-derived marker genes, such as *CYP11B2*[25] defining zona glomerulosa, *CYP11B1*[20] – zona fasciculata, *CYP17A1*, *SULT2A1*[26], and *CYB5A*[27] – zona reticularis. These genes were detected in all sample groups regardless of the extraction method, except for *CYP11B2*, which was not detected in the enzymatic and nuclei samples (**Figure 4a, Additional file 1: Figure S8a**). Similar results were obtained for the adrenal medullary tumor (n=15) and PitNET samples (n=9) with 94.807 (89% chromaffin cells) and 41.418 (79.6% pituitary cells) integrated cells, respectively (**Fig. 3c**). Correspondingly, these cells expressed key marker genes, such as *CHGA*, *SYP*[28], *DBH*, *PNMT*[29] (**Fig. 4b, Additional file 1: Figure S8b**) and *POMC* (only in the ACME HS datasets), with the exception of *GH1* and *POU1F1* in the ACME HS datasets (**Fig. 4c, Additional file 1: Figure S8c**). Although thyroid follicular cells represented the majority (95.2%) of the 60.365 cells in thyroid gland samples (n=5), ACME HS-dissociated cells exhibited almost no expression of key markers such as *TG* and *TSHR*[30],[31] (**Fig. 4d, Additional file 1: Figure S8d**).

**Fig. 4:**
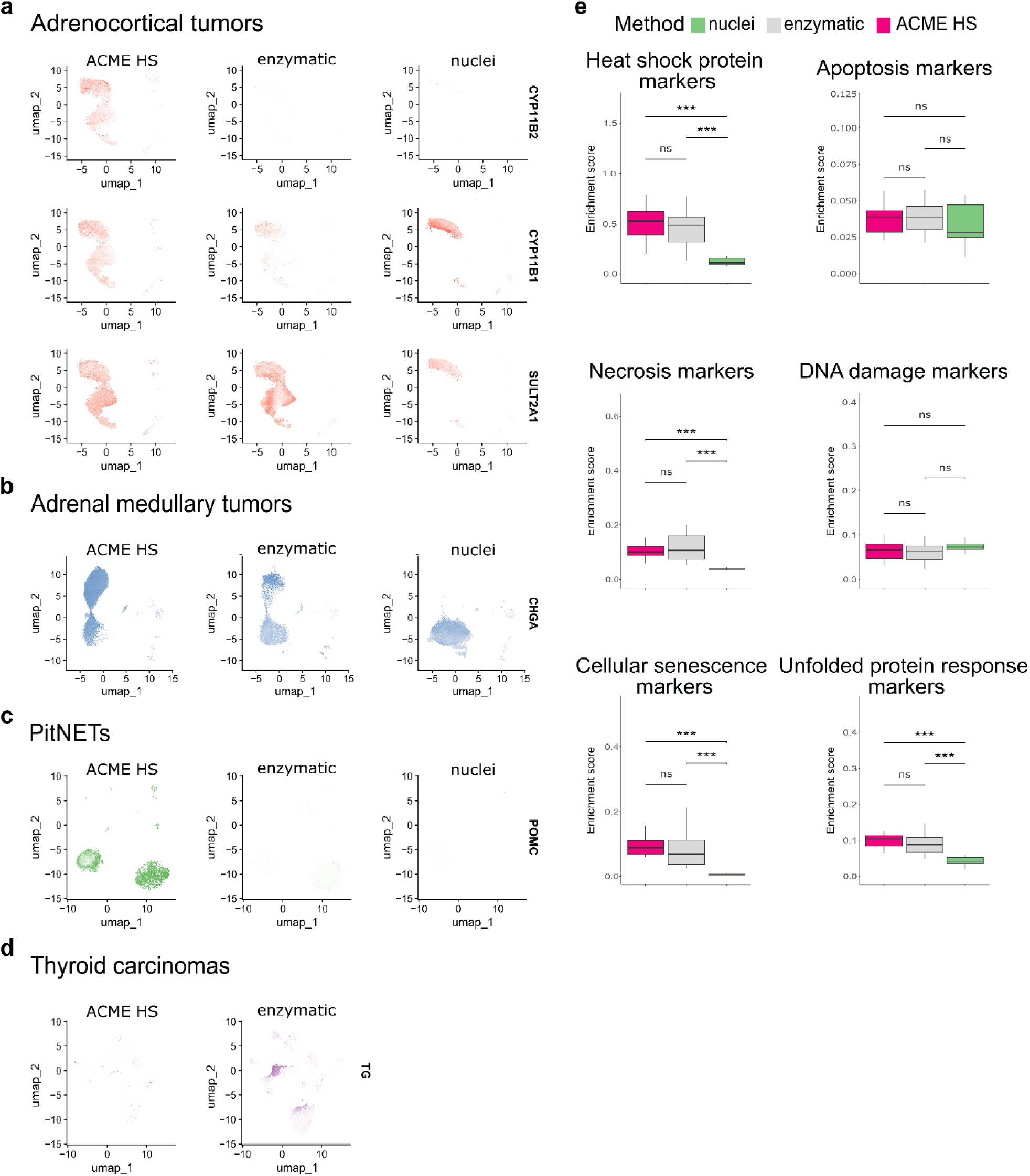
Expression of tissue-specific genes in nuclei and whole cells obtained by ACME HS and enzymatic dissociation methods. **a.** Key tissue-specific gene expression according to UMAP visualization namely, *CYP11B2, CYP11B1*, and *SULT2A1* for adrenocortical tumors; **b** *CHGA* for adrenal medullary tumors; **c** *POMC* for PitNETs; **d -** *TG* for thyroid carcinoma samples. **e**. The distribution of the enrichment scores of heat shock, apoptosis, necrosis, DNA damage, cell senescence, and unfolded protein response signatures across preparation methods of the datasets obtained from adrenocortical tumor and adrenal medullary tumor, thyroid carcinoma, and PitNET samples (combined dataset, n=41).

Then, we analysed a panel of the top genes specific for adrenocortical, chromaffin, thyroid follicular, and pituitary cells (**Additional file 1: Figure 8e**). It turns out that each dissociation method allows to estimate the expression signatures of different genes that have little overlap in ACME HS, enzymatic, and nuclear samples, with the exception of PitNETs.

### Difference in the expression of stress- and apoptosis-associated genes in the ACME HS, enzyme, and nuclei datasets

Since dissociation and preservation techniques can induce cellular stress[9],[5], as evidenced by changes at the transcriptomic level, we examined the expression of key markers of stress and cell death. Specifically, markers associated with apoptosis, necrosis, cellular senescence, DNA damage, heat shock, and the unfolded protein response (UPR) were assessed in the ACME HS (n=13), enzyme (n=19), and nuclei (n=9) datasets for all tissue samples (**Fig. 4e, Additional file 1: Figure S9, Additional files 2,3: Table S1,2**).

We found no differences in the expression levels of genes related to apoptosis in any of the datasets. The necrosis gene signature showed no differences between the ACME-HS and enzymatic samples but was lower in the nuclei. Nevertheless, the most significant expression of *TNF* (tumor necrosis factor) was observed in adrenocortical tumors and thyroid carcinomas in ACME HS and nuclei samples (**Additional file 3: Table S2)**. The DNA damage gene signature exhibited minimal variations among all three methods. In addition, the heat shock protein signature showed minimal differences between the ACME HS and enzymatic samples but was significantly lower for nuclei. The UPR signature did not differ between ACME HS and enzymatic samples but was significantly lower for nuclei. In particular, *ERN2,* a UPR marker, was highly expressed in adrenal medullary tumor and PitNET datasets obtained by ACME HS and nuclei isolation methods, in adrenocortical tumors by enzymatic digestion, and in thyroid carcinomas by the ACME HS method. In addition, the cellular senescence signatures also showed minimal differences between the ACME HS and enzymatic samples. *IL1B* was highly expressed in adrenocortical and medullary tumor samples obtained from the ACME HS and nuclei datasets, while *IL6* was highly expressed in all tissues compared with the PitNET datasets obtained by the enzymatic digestion method. Another important senescence marker, *CDKN2B,* was highly expressed in adrenal medullary tumor samples obtained by nuclei isolation as well as in thyroid carcinoma samples obtained by an enzymatic approach. Senescence markers *HMGA1* and *UBB* were highly expressed in the nuclei datasets for adrenocortical and medullary tumor samples (**Additional file 2: Table S2)**.

### RNA velocity estimates of the individual cells accurately recapitulated the transcriptional dynamics in the ACME HS and enzyme datasets

Next, we performed velocity analysis for individual samples to demonstrate the consistency between the ACME HS (n=1) and enzymatic (n=1) protocols together with their advantages over the nuclei isolation method (n=1). For adrenocortical (**Fig. 5a**) and chromaffin cells (**Fig. 5b**), we identified similar velocity directionalities from ACME HS and enzymatic-specific clusters towards cell populations commonly shared between methods. The RNA velocity recapitulated the transcriptional dynamics within these datasets, including the general movement of the differentiating adrenocortical and chromaffin cells, as well as movement towards and away from the intermediate differentiation state. The velocity also captured the cell cycle dynamics involved in cell differentiation.

**Fig. 5:**
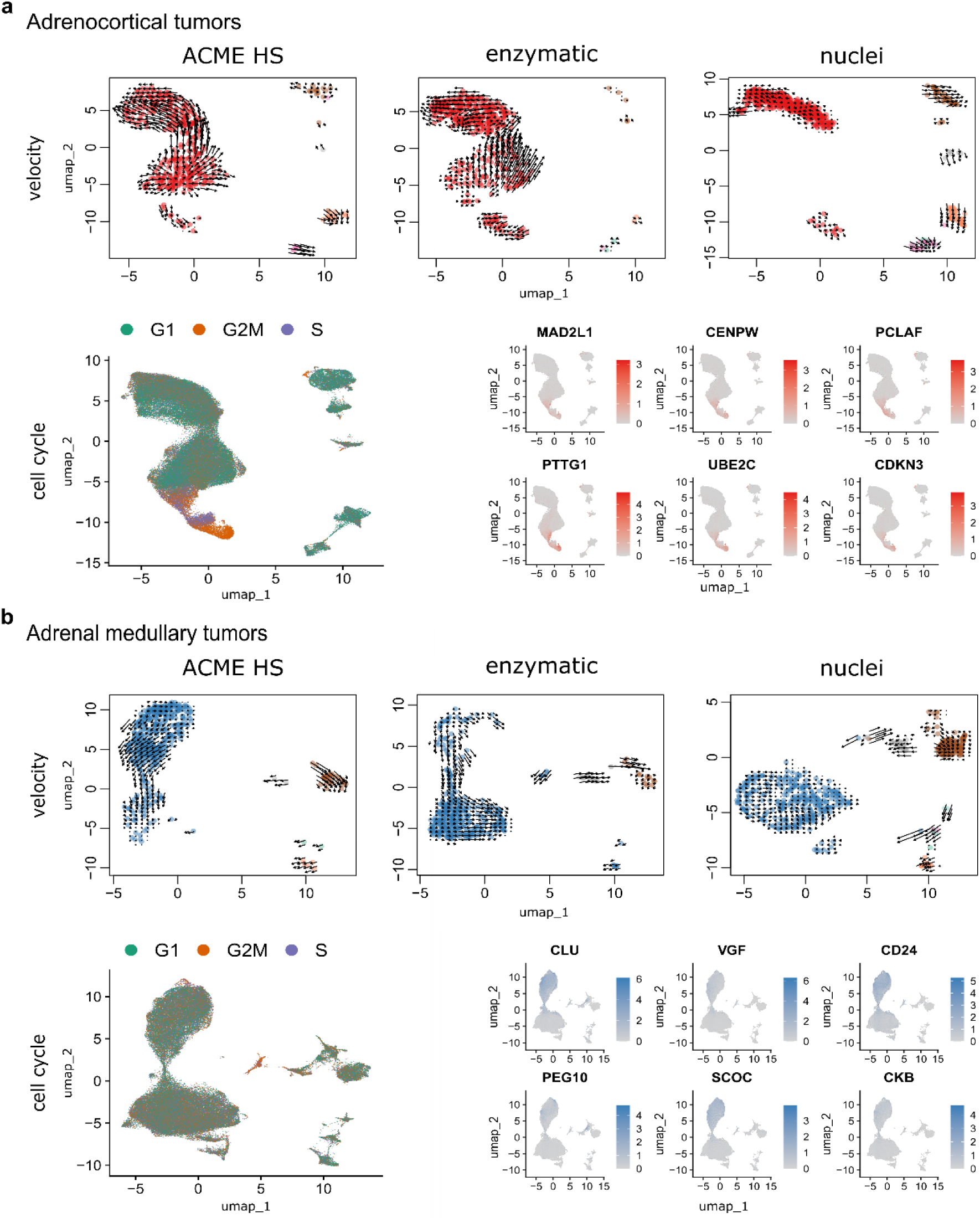
Velocity and cell cycle of the ACME HS, enzymatic and nuclei datasets. **a, b.** Velocity and cell cycle estimation for adrenocortical and adrenal medullary tumor datasets, respectively. Velocity was performed for individual samples (n=1) for each method, and cell cycle estimation was performed for adrenocortical (n=12) and adrenal medullary tumors (n=15). Examples of differentially expressed (DE) genes associated with cell cycle control are shown on the individual embeddings. DE analysis was conducted for specific clusters within major cell types, such as A-7 and C-4 (**Additional file 1: Figure S6**).

We observed G2M and S phase cells at velocity start point in ACME HS and enzymatic specific clusters as well as differential expression of cell cycle controlling and neuroendocrine tumor proliferation genes – *MAD2L1, CENPW, PCLAF, PTTG1, UBE2C*, *CDKN3* for adrenocortical cells and *CLU, VGF, CD24, PEG10, SCOC*, *CKB* for chromaffin cells in all samples. We suggest that cells expressing these genes are potential adrenocortical and chromaffin cell progenitors, respectively (**Fig. 5a, b**). Thus, the ACME HS method is suitable for studying intermediate differentiation states of cancer and progenitor cell populations. Although proliferating cells in PitNETs were determined as a separate cell cluster, we did not find a clear-cut pattern in RNA velocity and the cell cycle (**Additional file 1: Figure S10**).

## Discussion

In this study, we present an optimized ACME HS[17] dissociation technique for the effective isolation of single cells from flash-frozen human tissues. In the real-life setting of clinical research centers, the utilization of fresh tissues is rather complicated and frequently disruptive for cells that are to be further analysed using scRNA-seq (see *Introduction for details*). Consequently, the most frequent alternative is the use of fresh-frozen tissues obtained during biobanking as a starting material. However, mechanical and physical stress inflicted upon cells during freeze-thawing invariably results in a degree of cell membranes rupture[32], resulting in the release of large amounts of freely floating transcripts (ambient RNA) into cell suspensions, thereby contaminating endogenous gene expression profiles and confounding cell type annotation results.

In contrast, the ACME HS technique starts with frozen material, thereby overcoming the limitations associated with the freeze-thawing of living cells. Probably, even more importantly, simultaneous acetic acid-based dissociation and methanol-based fixation “snaps” the transcriptional profiles of individual cells at the very beginning of the procedure, thereby eliminating the global transcriptome changes associated with the action of dissociation enzymes and displacement of cells from their original tissue context/microenvironment. Yet, another important point is that the ACME HS allows for recovery of high-integrity RNA even following cryopreservation of fixed cell suspensions. Finally, in contrast to single nuclei isolation techniques, ACME HS preserves the cytoplasm of cells, yielding a dramatically better representation of mature mRNAs.

Here, we demonstrated that ACME HS is suitable and efficient technique for isolating of the high-quality single cell suspensions from difficult-to-dissociate complex tissues with high lipid content and large areas of fibrosis and calcinosis, the tissues typically posing significant challenges for enzymatic digestion. We have successfully obtained 304 465 cells from 41 endocrine neoplasms, namely, adrenal medullary tumors, adrenocortical tumors, thyroid follicular cell-derived carcinomas and the pituitary-derived neuroendocrine tumors by ACME HS, enzymatic, and nuclei isolation methods.

One of the key optimization points as compared to the original ACME technique was the use of the high-salt 3xSSC washing buffer instead of PBS. The main rationale behind this point is that, under physiological ionic strength, RNAses may be reactivated during the rehydration, thereby dramatically diminishing the yield and integrity of mRNA and ribosomes. In that, 3xSSC supplemented with DTT and RNase inhibitor dehydrates the cells and blocks the activity of RNAses[33], allowing for an efficient preservation of nucleic acids inside the cells.

Using flow cytometry, we were able to provide evidence for obtaining of sufficient cell numbers employing ACME HS technique, with standard DNA histograms and accurately determined cell cycle phases further confirming the proper processing of our samples. The degree of cellular debris and subG1-fragments may be attributed to the freeze-thawing of the samples during both the enzymatic and ACME HS protocols and does not compromise our conclusions.

We further assessed the cellular stress responses associated with the different sample processing techniques. Since the stress response genes are known to be activated upon the proteolytic tissue dissociation at 37°C, we expected the major differences in the expression profiles there of between ACME HS/single nuclei isolation performed under the ice-cold conditions vs enzymatic dissociation protocol performed at 37°C. However, despite our initial considerations, the total contributions of the stress signatures (heat shock, necrosis, cellular senescence, and UPR) in enzymatic and ACME HS dissociation protocols of tumor tissues were essentially the same, with exposure to collagenase and membrane rupture during methanol incubation (causing loss of cytoplasmic mRNA) being deduced as the major stress factors in enzymatic and ACME HS protocols, respectively. The nuclei isolation protocol significantly outperformed both enzymatic and the ACME HS dissociation methods in terms of the reduced stress responses identified in scRNA/snRNA-Seq profiles. Both protocols for the isolation of single cell suspensions performed significantly better than isolation of single nuclei in the majority of the other comparisons in our study. However, this specific issue may be highly relevant in studies, where minimizing of sample processing-associated stress responses and/or enrichment of sequencing data with intronic sequences in nuclear immature RNAs are of critical importance, dictating the choose of isolation of single nuclei instead of single cells’ isolation in these cases.

Going to the whole-transcriptome level, we were able to successfully integrate the ACME HS, enzyme, and nuclei datasets, further integrating them with the reference scRNA-Seq profiles of the cognate normal tissues. We examined all of the acquired datasets (ACME HS, enzymatic, nuclei) to evaluate the heterogeneity of the major cell populations (adrenocortical, chromaffin, thyroid follicular, and pituitary neuroendocrine cells) for all four tissues studied. Overall, we demonstrated a comparable representation of the major cell types, subpopulations, and functional states in ACME HS and enzymatic methods, while the single nuclei-based protocol performed significantly much worse. Our data thus corroborate previous observations on the principal differences of scRNA vs snRNA profiles, particularly in terms of cytoplasm-associated signatures, including those associated with cellular metabolism[34], protein synthesis and mRNA processing[35],[36], the processes being particularly important for tumorigenesis[37],[38], with a proper representation thereof being critical for obtaining of the biologically relevant data in the studies of human neoplastic diseases.

Finally, we performed the velocity analysis and assessment of the activity of cell cycle markers to demonstrate that nuclei-based data were largely depleted from the information on putative differentiation directions, intron-retention events, as well as cell cycle phases connectivity. Again, the data obtained from both single cell dissociation protocols were fairly consistent, implying thereof as preferable approaches for studying cell differentiation, clonal evolution in cancer and intron-retention events.

In summary, we optimized and employed the ACME HS technique for the scRNA analysis of human tissues derived from various endocrine neoplasms. We clearly demonstrated that scRNA profiling of single cell suspensions obtained using ACME HS and enzymatic methods significantly outperformed snRNA profiling in terms of marker gene expression analysis and tumorigenesis while demonstrating in-between comparable performances in the majority of implemented analyses. Additionally, the modified ACME protocol allows for an extra-option of successful cryopreservation of dissociated/fixed cells without sacrificing the mRNA yield and integrity. To our knowledge, this is the first report on successful implementation of the ACME HS technique in primary human tissues, and we believe that this protocol should significantly promote the scRNA studies in humans that are to be explicitly compliant with the real-life infrastructure and logistics of the surgical care centers.

## Conclusions

The key determining factor for a successful scRNA-seq profiling is a robust sample preparation step that yields a high-quality single-cell suspension. In our study, we adapted the ACME HS dissociation approach for fresh-frozen endocrine tissues. We demonstrated that ACME HS dissociates and fixes cells with preserved morphology and high RNA integrity number and maintains the cytoplasm of the cells, yielding a better representation of the mature mRNAs. We obtained 304 465 cells and nuclei and demonstrated a comparable representation of the major cell types, subpopulations, and functional states in ACME HS and enzymatic dissociation methods. We showed that scRNA analysis of single cells obtained by ACME HS and enzymatic methods significantly outperformed snRNA analysis regarding marker gene expression and velocity analysis. We showed that ACME HS is the first practical approach for obtaining fixed intact cells from fresh-frozen tissues for scRNA-seq studies of human tissues.

## Methods

### Tissue sampling

Fifty-three human adrenal gland neoplasms (including 17 adrenocortical tumors and 36 adrenal medullary tumors), 12 thyroid carcinomas, and 38 pituitary neuroendocrine tumors (PitNETs) were acquired from the Endocrinology Research Centre, Moscow, Russia (**Additional file 2: Table S1**). In all patients, tumor specimens were definitively diagnosed by imaging, surgery, and histopathological examination. Each study participant gave written informed consent. In addition, single cells were isolated from fresh and fresh-frozen adrenal medullary tumor, adrenocortical tumor, thyroid carcinoma, and PitNET samples. After sampling, the tissues were placed in a cold Tissue Storage Solution (Miltenyi Biotec) pending dissociation. The fresh-frozen samples were stored at - 80°C until processed.

### ACME HS dissociation

After tissue sampling, 200-250 mg of fresh-frozen adrenal gland neoplasms, thyroid carcinoma, or 5-10 mg PitNET samples were thoroughly minced on ice and immediately added to ACME solution (15% methanol, 0.1M glacial acetic acid, 0.1M glycerol, 0.1M N-acetyl cysteine (NAC) and RNase-free water) to achieve a volume of 10 ml in 15 ml Falcon tube. NAC cleans cells from mucus, fatty lipids and protects cells from oxidative damage. The samples were dissociated at room temperature for 1 h on a shaker set at 35 rpm with vertical platform rotation. During incubation, the mixture was carefully pipetted 2-4 times using 5 ml pipette tips. After incubation, the samples were centrifuged at 1000xg for 5 minutes at 4°C to remove the ACME solution. From this point, the samples were kept on ice. The supernatant was carefully discarded, and 2-4 ml of cold 3xSSC* buffer (3xSSC, 40 mM DTT, 1% BSA, and RNase-free water) containing 0.5 U/µL of the RNase Inhibitor RiboLock (Thermo Fisher Scientific) was added to the cell pellet and resuspended.

The homogenate was sequentially filtered through a pre-wetted (with 500uL 3xSSC*) 70μm and 40μm filters (Miltenyi Biotec) into a 15 ml tube, and centrifuged at 1000xg for 7 minutes (4 °C). The supernatant was then carefully removed, and the pellet was resuspended in 1-2 ml of cold 3xSSC*.

### Enzymatic dissociation

Approximately 200-250 mg of fresh adrenal gland neoplasm, thyroid carcinoma, or 5-10 mg of PitNET samples were washed in HBSS, thoroughly minced on ice, and placed in dissociating solution at 37°C with gentle pipetting every 5 minutes.

Adrenal gland neoplasm samples were dissociated with 25-30 ul of enzyme D Multi Tissue Dissociation Kit (MTDK) (Miltenyi Biotec) or enzyme A Neural Tissue Dissociation Kit P (NTDK) (Miltenyi Biotec) or 2 mg/ml collagenase IV (Gibco, Thermo Fisher Scientific) or collagenase I (Gibco, Thermo Fisher Scientific) in 870 mM HBSS, 10% FBS, and 20 mM HEPES for 20-30 minutes.

Thyroid carcinoma samples were dissociated with 30-35 ul of enzyme D Multi Tissue Dissociation Kit (Miltenyi Biotec) or 2 mg/ml collagenase IV (Gibco, Thermo Fisher Scientific) in 870 mM HBSS, 10% FBS, and 20 mM HEPES for 20-30 minutes.

PitNET samples were dissociated with 8-10 ul of enzyme D Multi Tissue Dissociation Kit (Miltenyi Biotec) or 2 mg/ml collagenase IV (Gibco, Thermo Fisher Scientific) in 870 mM HBSS, 10% FBS, and 20 mM HEPES for 7-15 minutes.

The obtained homogenate was filtered in 3-5 ml of Wash Buffer (1x DPBS, containing 10% FBS, 20mM HEPES, and 6mM glucose) through a prewetted 70μm cell culture filter (Miltenyi Biotec) and centrifuged for 5 minutes at 300xg (4 °C).

For samples with high blood and debris content, the red blood cells were lysed using Red Blood Cell Lysis Solution (Miltenyi Biotec), and dead cells were removed with Dead Cell Removal Kit (Miltenyi Biotec). The cells were counted and assessed for viability using trypan blue staining on Countess 3 (Thermo Scientific). After all, the pellet was resuspended in a Wash Buffer volume of 100-400 ul, depending on the pellet size.

### Nuclei isolation

Nuclei were isolated from fresh-frozen adrenocortical tumor, adrenal medullary tumor, and PitNET specimens. Fresh-frozen tissue samples were thoroughly minced on ice and placed into a gentleMACS C tube (Miltenyi Biotec) with 2ml ice-cold Hypotonic lysis buffer (10mM HEPES pH7.2, 5mM MgCl2, 10mM NaCl, and 1% NP40). GentleMACS C tubes were then placed on the gentleMACS Dissociator (Miltenyi Biotec) and the samples were homogenized by running the program h_mito_01, and then incubated on ice for 10 minutes. After repeating the homogenization step, 2 ml of Isotonic buffer (10mM HEPES pH7.2, 5mM MgCl2,10mM NaCl, and 500mM sucrose) was added to the lysates, mixed by pipetting, filtered through a pre-wetted 70μm cell culture filter (Miltenyi Biotec) with Isotonic wash buffer (10mM HEPES pH7.2, 5mM MgCl2,10mM NaCl, and 250mM sucrose) and centrifuged for 5 minutes at 1000g (4 °C). Then, we carefully removed the supernatant, resuspended the pellet in 1 ml of DPBS with 1% BSA, and filtered it through a prewetted 30μm cell culture filter (Miltenyi Biotec). After all, the pellet was resuspended in DPBS containing 1% BSA volume 100-400 ul, depending on the pellet size.

### RNA extraction and quality assessment

To evaluate the RIN of the samples depending on the duration of their storage, we isolated RNA from cell suspensions prepared by the ACME HS and enzymatic dissociation methods. We isolated RNA from fresh/fresh-frozen ACME HS and enzyme-dissociated cells after dissociation (0 days) and interval cryopreservation or freezing (1, 3, 7, 14, and 28 days). RNA extractions were performed using an AllPrep DNA/RNA Mini Kit (QIAGEN), following the manufacturer’s protocol. RNA quality was assessed using an Agilent 5200 Fragment Analyzer, using the Agilent HS RNA (15NT) kit.

### Immunocytochemistry

Immunocytochemistry was performed for the markers CYP11B1 and TSHR to identify the nature of adrenocortical and thyroid cells. Initially, membranes of enzyme-dissociated cells were permeabilized with 100 ul of permeabilization enzyme (10x genomics, 2000214) for 20 minutes at 37°C. After that, the ACME HS and enzyme-dissociated cells were blocked in 3% BSA for 20 minutes. The cells were incubated in the diluent buffer (ab64211) of the primary polyclonal Anti-CYP11B1 antibody (ab197908) and the TSH Receptor monoclonal antibody (4C1) (Invitrogen, MA5-16519) at a dilution of 1:200 for 1 h at 4°C. The cells were washed in antibody diluent buffer before being incubated for 1 h at 4°C with the secondary antibody AlexaFluor 594 (ab150080) or AlexaFluor 594 (ab150116) at a dilution of 1:500, respectively. After repeating the washing step, the cells were stained with Hoechst 33342 (BD Pharmingen™). Visualization of the antigen-antibody complexes was performed using an Olympus FV3000 Scanning Confocal Microscope (Olympus corporation, Tokyo, Japan).

### Methanol fixation and ACME HS cryopreservation

For methanol fixation, we took 200 ul of previously prepared enzyme-dissociated cells in wash buffer (*Methods, Enzymatic dissociation*) with 0.5 U/µL of the RNase Inhibitor RiboLock (Thermo Fisher Scientific) and added 800 ul of 100% ice-cold methanol drop by drop to the cells while gently vortexing the tube to avoid clumping of the cells. We stored the fixed cells at −80°C.

For ACME HS cryopreservation, we took 900 ul of cell suspension in 3xSSC*(*Methods, ACME HS*) and cryopreserved them with 10% DMSO. Store the fixed cells at −80°C.

### Flow cytometry

ACME HS and enzyme-dissociated cells isolated from adrenal gland neoplasm, thyroid carcinoma, and PitNET samples were transferred into DPBS with 0.1% FBS at a final concentration of 106 cells/ml. Next, 2 µM of 5.6-carboxyfluorescein diacetate succinimidyl ester or CFSE (BD Biosciences, USA) was added to the cells and incubated for 5 minutes at 370C. Then, the cells were washed twice with 10 volumes of cold DPBS, and stained with PI (10 µg/ml) in 0.5 ml of PI/RNase staining buffer (BD Biosciences, USA) for 30 minutes at room temperature in the dark.

Flow cytometric analysis was performed on a NovoCyte 2060R machine (Agilent, USA) equipped with two lasers, including a laser tuned at 488 nm to excite CFSE and PI, and the standard set of detectors for green fluorescence of CFSE and red fluorescence of PI. Program compensation was used to correct spectral spillover. Fluidics and optics were calibrated with NovoCyte QC particles. The threshold was set at FSC-H. Samples were run at the lowest flow rate. At least 10,000 events were analysed. Deconvolution of the DNA histograms was performed with the instrument Software NovoExpress.

### Histopathological examination

Tumor tissue samples obtained during surgical treatment of patients at the Endocrinology Research Center were fixed in 10% buffered formalin, processed in the histological staining system of a Leica ASP6025, and embedded in paraffin. Subsequently, paraffin sections with a thickness of 3 µm were cut from the paraffin-embedded tumor tissue samples using a microtome and applied to slides treated with poly (l-lysine). The slides were then stained with hematoxylin and eosin following the standard procedure. All histological slides were scanned using a Leica Aperio AT2 system at 20x magnification for further analysis.

### Preparation of the cell suspensions for loading on the 10x Chromium controller

ACME HS-cryopreserved and methanol-fixed cells were unfrozen and centrifuged at 2000x g for 5 minutes (4 °C) to remove the 3xSSC*/DMSO and methanol. After that, the pellet was resuspended in cold 3xSSC* buffer to a density of some 2000 cells or nuclei/ul.

### scRNA-seq and snRNA-seq using the 10X Genomics platform

Single cells or nuclei were captured and barcoded, and cDNA libraries were generated using the Chromium Next GEM Single Cell 3ʹGEM, Library & Gel Bead Kit v3.1 (10X Genomics). For each sample, 10 000 cells or nuclei (∼ 2000 cells or nuclei in 1ul, cell suspension volume calculator table 10X Genomics) in cold 3xSSC* were mixed with RT-PCR master mix and immediately loaded together with Single-Cell 3′ Gel Beads and Partitioning Oil into a Chromium Chip G. cDNA, and gene expression libraries were generated according to the manufacturer’s instructions (10x Genomics). cDNA and gene expression libraries were quantified using a Qubit dsDNA HS assay kit (Thermo Fisher Scientific), and the cDNA and gene expression library fragment sizes were assessed with the Agilent 5200 Fragment Analyzer, using a DNA HS (1-6000NT) kit. The final libraries were multiplexed and sequenced on an Illumina Novaseq 6000 platform, using the S4 Reagent Kit v1.5 (200 cycles).

### Adrenal, thyroid and pituitary glands single-cell transcriptomic analysis

The Raw sequenced reads were processed with 10X Cell Ranger (v6.1.1). Default Cell Ranger quality check measurements were used for further comparison methods through the Wilcoxon test. The expression matrixes for the filtered cells were submitted to Seurat[24] (v4.9.9 and v5.0.0) for basic analysis, including scaling and normalization. Cell filtering based on gene/molecule dependency was done by pagoda2[39] (v1.0.11). Doublets and ambient RNA content were calculated with scrublet[40] (v0.2.3), SoupX[41] (v1.6.2) and decontX[42] (v3.18), respectively, with default settings. The means for doublets and ambient RNA values per sample were compared between sample preparation methods (ACME HS, enzymatic, nuclei). Major cell types were identified by the label propagation function using Conos[43] (v1.5.0) and reference datasets[44],[45],[46],[47],[48]. Sample integration was conducted by applying RunHarmony on preprocessed Seurat objects. Velocity analysis was performed with Velocyto (v0.17) on 1000 cells subset per sample and visualized using Velocyto.R[49] (v0.6) on integrated embeddings. Cell cycle phase predictions were based on reference gene expression processed with Seurat (v5.0.0). Differential expression was conducted for all cells of different sample preparation methods. A functional enrichment test was performed for differentially expressed genes with clusterProfiler[50] and wiki pathways as reference databases.

### Stress and cell death gene signatures

We utilized the PercentageFeatureSet function with default parameters from the Seurat package to evaluate the impact of various methods - ACME HS, enzymatic dissociation, and nuclei isolation on cellular stress, focusing on determining their potential to induce stress or favored necrosis or apoptosis among the cells. This function computes the percentage of all counts assigned to a specified set of genes. For the apoptosis signature, we curated a gene signature encompassing *CASP3, BAX, BAD, BID, APAF1, TP53, FAS, TNFRSF10B, CYCS, BCL2,* and *AIFM1*[51]. Meanwhile, for the necrosis signature, we selected *HMGB1*[52]*, ATP5F1A, CALR, ARHGAP45, S100A8, S100A9, NAMPT, ANXA1, KRT18, TNF,* and *AGER* genes[53],[54]. We assessed various modalities of cell stress, including oxidative stress, cellular senescence, DNA damage, heat shock, and the unfolded protein response. The oxidative stress signature was constructed using *NFE2L2, KEAP1, SOD1, CAT, HMOX1, GCLC, GCLM, NQO1,* and *PRDX1* genes[53],[54]. The markers of cellular senescence included *CDKN1A, CDKN2A, IGFBP3, GADD45A, CCND1, CDKN2B, IL1A, IL1B, IL6, IL10, HMGA1, HMGB2,* and *UBB*[53],[54]. DNA damage signatures comprised *TP53, BRCA1, CHEK2, ATM, RAD51, RPA1, MDM2, ATR,* and *XRCC5*[53],[54]. For the heat shock signature, we considered the HSP family genes *HSPB, HSPG2, HSPB11, HSPA6, HSPD1, HSPE1, HSPBAP1, HSPA4L, HSPB3, HSPA4, HSPA9, HSPA1L, HSPA1A, HSPA1B, HSP90AB1, HSPB1, HSPA5, HSPA14, HSPA14.1, HSPA12A, HSPB2, HSPA8, HSP90B1, HSPB8, HSPH1, HSPA2, HSP90AA1, HSPB9, HSPB6, HSPBP1, HSPA12B,* and *HSPA13*[55],[56]. Lastly, the unfolded protein response signature included *ATF4, ATF6, XBP1, HSPA5, DDIT3, HERPUD1, DNAJC3, ERN1, ERN2,* and *PDIA6* genes[53],[54]. Using a two-tailed Wilcoxon rank-sum test, we calculated statistically significant differences in the signature scores between the various dissociation methods.

### Statistical Data Analysis

All the data were presented as the means and standard deviations. Statistical significance (assessed by two-tailed t-test and Wilcoxon rank-sum test) is shown in the figures: **** (0.0001 < *p* < 0.001), *** (*p* < 0.001), ** (0.001 < *p* < 0.01), * (0.01 < *p* < 0.05), ns - not significant - *p* > 0.05.

## Declarations

### Ethics approval and consent to participate

The research was performed in accordance with the Declaration of Helsinki. The studies involving human participants were reviewed and approved by the local Ethics Committee of the Endocrinology Research Centre (Protocol No. 16 dated 14.10.2020). Written informed consent to participate in this study was provided by all patients.

### Consent for publication

Not applicable

### Availability of data and materials

All scRNA-seq datasets generated and analysed during this current study have been deposited in the Gene Expression Omnibus (GEO) database under accession codes: GSEXXXXX (AdrenocorticalTumors_ACME), GSEXXXXX (AdrenocorticalTumors_enzymatic), GSEXXXXX (AdrenocorticalTumors_nuclei), GSEXXXXX (MedullaryTumors_ACME), GSEXXXXX (MedullaryTumors_enzymatic), GSEXXXXX (MedullaryTumors_nuclei), GSEXXXXX (PitNETs_ACME), GSEXXXXX (PitNETs_enzymatic), GSEXXXXX (PitNETs_nuclei), GSEXXXXX (ThyroidCarcinomas _ACME), GSEXXXXX (ThyroidCarcinomas_enzymatic).

### Author Contributions

**MU** and **AS** curated the data, designed and performed numerous experiments, analysed the data, and drafted the manuscript. **RD**, **VT, DM** and **EA** performed the data analysis. **AR** conceptualized and supported setting up of ACME HS protocols for cell preservation. **MYL** and **MP** performed flow cytometry. **AK**, **AG** and **WA** collected and sequenced endocrine tumor samples. **RS** and **LU** made microscopic images. **DB, LU**, **EB**, **AL,** and **AS** provided and performed the collection of human clinical materials. **SP** and **OG** supervised, coordinated and conceptualized the work. **LD, IM**, **GM**, **ID,** and **NM** managed responsibility for the research activity planning and execution. All authors read and approved the final manuscript.

## Acknowledgements

We thank the Center for Personalized Medicine of Endocrine Diseases members for their valuable comments and critical discussion. We thank Dr. Pavel Volchkov at Lomonosov Moscow State University (Moscow, Russia) for assembling a proficient cohort of individuals who share a common scientific passion, resulting in a successful team. We also thank Julia Krupinova at Moscow Clinical Scientific Center N.A. A.S. Loginov (Moscow, Russia) for supporting the service design documentation. We thank Ekaterina Markelova at the Endocrinology Research Centre (Moscow, Russia) for the storage and cryopreservation of tumor samples.

## Reference

1. Zhang S, Cui Y, Ma X, Yong J, Yan L, Yang M, et al. Single-cell transcriptomics identifies divergent developmental lineage trajectories during human pituitary development. Nat Commun. 2020;11:5275.

2. Kameneva P, Melnikova VI, Kastriti ME, Kurtova A, Kryukov E, Murtazina A, et al. Serotonin limits generation of chromaffin cells during adrenal organ development. Nat Commun. 2022;13:2901.

3. Dong R, Yang R, Zhan Y, Lai H-D, Ye C-J, Yao X-Y, et al. Single-Cell Characterization of Malignant Phenotypes and Developmental Trajectories of Adrenal Neuroblastoma. Cancer Cell. 2020;38:716–733.e6.

4. Slyper M, Porter CBM, Ashenberg O, Waldman J, Drokhlyansky E, Wakiro I, et al. A single-cell and single-nucleus RNA-Seq toolbox for fresh and frozen human tumors. Nat Med. 2020;26:792– 802.

5. van den Brink SC, Sage F, Vértesy Á, Spanjaard B, Peterson-Maduro J, Baron CS, et al. Single-cell sequencing reveals dissociation-induced gene expression in tissue subpopulations. Nat Methods. 2017;14:935–6.

6. Waymouth C. To disaggregate or not to disaggregate injury and cell disaggregation, transient or permanent? In Vitro. 1974;10:97–111.

7. Cerra R, Zarbo RJ, Crissman JD. Dissociation of cells from solid tumors. Methods Cell Biol. 1990;33:1–12.

8. Cunningham RE. Tissue disaggregation. Methods Mol Biol. 2010;588:327–30.

9. Denisenko E, Guo BB, Jones M, Hou R, de Kock L, Lassmann T, et al. Systematic assessment of tissue dissociation and storage biases in single-cell and single-nucleus RNA-seq workflows. Genome Biol. 2020;21:130.

10. Lake BB, Ai R, Kaeser GE, Salathia NS, Yung YC, Liu R, et al. Neuronal subtypes and diversity revealed by single-nucleus RNA sequencing of the human brain. Science. 2016;352:1586–90.

11. O’Flanagan CH, Campbell KR, Zhang AW, Kabeer F, Lim JLP, Biele J, et al. Dissociation of solid tumor tissues with cold active protease for single-cell RNA-seq minimizes conserved collagenase-associated stress responses. Genome Biology. 2019;20:210.

12. Deleersnijder D, Callemeyn J, Arijs I, Naesens M, Van Craenenbroeck AH, Lambrechts D, et al. Current Methodological Challenges of Single-Cell and Single-Nucleus RNA-Sequencing in Glomerular Diseases. J Am Soc Nephrol. 2021;32:1838–52.

13. Schmøkel SS, Nordentoft I, Lindskrog SV, Lamy P, Knudsen M, Jensen JB, et al. Improved protocol for single-nucleus RNA-sequencing of frozen human bladder tumor biopsies. Nucleus. 2023;14:2186686.

14. Thrupp N, Sala Frigerio C, Wolfs L, Skene NG, Fattorelli N, Poovathingal S, et al. Single-Nucleus RNA-Seq Is Not Suitable for Detection of Microglial Activation Genes in Humans. Cell Rep. 2020;32:108189.

15. Massoni-Badosa R, Iacono G, Moutinho C, Kulis M, Palau N, Marchese D, et al. Sampling time-dependent artifacts in single-cell genomics studies. Genome Biol. 2020;21:112.

16. Adam M, Potter AS, Potter SS. Psychrophilic proteases dramatically reduce single-cell RNA-seq artifacts: a molecular atlas of kidney development. Development. 2017;144:3625–32.

17. García-Castro H, Kenny NJ, Iglesias M, Álvarez-Campos P, Mason V, Elek A, et al. ACME dissociation: a versatile cell fixation-dissociation method for single-cell transcriptomics. Genome Biology. 2021;22:89.

18. Canmillo Schneider K. Histologie von Hydra fusca mit besonderer Berücksichtigung des Nervensystems der Hydropolypen. 1890. https://link.springer.com/article/10.1007/bf02955882. Accessed 4 Oct 2023.

19. David CN. A quantitative method for maceration of hydra tissue. Wilhelm Roux Arch Entwickl Mech Org. 1973;171:259–68.

20. Duparc C, Camponova P, Roy M, Lefebvre H, Thomas M. Ectopic localization of CYP11B1 and CYP11B2-expressing cells in the normal human adrenal gland. PLoS One. 2022;17:e0279682.

21. Crowley LC, Marfell BJ, Waterhouse NJ. Analyzing Cell Death by Nuclear Staining with Hoechst 33342. Cold Spring Harb Protoc. 2016;2016.

22. Darzynkiewicz Z. Critical aspects in analysis of cellular DNA content. Curr Protoc Cytom. 2011;Chapter 7:7.2.1–7.2.8.

23. Dumitriu IE, Mohr W, Kolowos W, Kern P, Kalden JR, Herrmann M. 5,6-carboxyfluorescein diacetate succinimidyl ester-labeled apoptotic and necrotic as well as detergent-treated cells can be traced in composite cell samples. Anal Biochem. 2001;299:247–52.

24. Hao Y, Stuart T, Kowalski MH, Choudhary S, Hoffman P, Hartman A, et al. Dictionary learning for integrative, multimodal and scalable single-cell analysis. Nat Biotechnol. 2023;:1–12.

25. van de Wiel E, Chaman Baz A-H, Küsters B, Mukai K, van Bonzel L, van Erp M, et al. Changes of the CYP11B2 Expressing Zona Glomerulosa in Human Adrenals From Birth to 40 Years of Age. Hypertension. 2022;79:2565–72.

26. Janšáková K, Hill M, Čelárová D, Celušáková H, Repiská G, Bičíková M, et al. Alteration of the steroidogenesis in boys with autism spectrum disorders. Transl Psychiatry. 2020;10:340.

27. Nakamura Y, Xing Y, Hui X-G, Kurotaki Y, Ono K, Cohen T, et al. Human adrenal cells that express both 3β-hydroxysteroid dehydrogenase type 2 (HSD3B2) and cytochrome b5 (CYB5A) contribute to adrenal androstenedione production. J Steroid Biochem Mol Biol. 2011;123:122–6.

28. Mete O, Asa SL, Gill AJ, Kimura N, de Krijger RR, Tischler A. Overview of the 2022 WHO Classification of Paragangliomas and Pheochromocytomas. Endocr Pathol. 2022;33:90–114.

29. Konosu-Fukaya S, Omata K, Tezuka Y, Ono Y, Aoyama Y, Satoh F, et al. Catecholamine-Synthesizing Enzymes in Pheochromocytoma and Extraadrenal Paraganglioma. Endocr Pathol. 2018;29:302–9.

30. Lee HJ, Li CW, Hammerstad SS, Stefan M, Tomer Y. Immunogenetics of Autoimmune Thyroid Diseases: A comprehensive Review. J Autoimmun. 2015;64:82–90.

31. Liu Y, Liu C, Pan Y, Zhou J, Ju H, Zhang Y. Pyruvate carboxylase promotes malignant transformation of papillary thyroid carcinoma and reduces iodine uptake. Cell Death Discov. 2022;8:423.

32. Isolated nuclei from frozen tissue are the superior source for single cell RNA-seq compared with whole cells | bioRxiv. https://www.biorxiv.org/content/10.1101/2023.02.19.529150v1. Accessed 13 Feb 2024.

33. Chen J, Cheung F, Shi R, Zhou H, Lu W, CHI Consortium. PBMC fixation and processing for Chromium single-cell RNA sequencing. J Transl Med. 2018;16:198.

34. Gaedcke S, Sinning J, Dittrich-Breiholz O, Haller H, Soerensen-Zender I, Liao CM, et al. Single cell versus single nucleus: transcriptome differences in the murine kidney after ischemia-reperfusion injury. Am J Physiol Renal Physiol. 2022;323:F171–81.

35. Santiago CP, Gimmen MY, Lu Y, McNally MM, Duncan LH, Creamer TJ, et al. Comparative Analysis of Single-cell and Single-nucleus RNA-sequencing in a Rabbit Model of Retinal Detachment-related Proliferative Vitreoretinopathy. Ophthalmol Sci. 2023;3:100335.

36. Bakken TE, Hodge RD, Miller JA, Yao Z, Nguyen TN, Aevermann B, et al. Single-nucleus and single-cell transcriptomes compared in matched cortical cell types. PLoS One. 2018;13:e0209648.

37. Venit T, Sapkota O, Abdrabou WS, Loganathan P, Pasricha R, Mahmood SR, et al. Positive regulation of oxidative phosphorylation by nuclear myosin 1 protects cells from metabolic reprogramming and tumorigenesis in mice. Nat Commun. 2023;14:6328.

38. Zhang Y, Qian J, Gu C, Yang Y. Alternative splicing and cancer: a systematic review. Signal Transduct Target Ther. 2021;6:78.

39. pagoda2. 2023.

40. Wolock SL, Lopez R, Klein AM. Scrublet: Computational Identification of Cell Doublets in Single-Cell Transcriptomic Data. Cell Syst. 2019;8:281–291.e9.

41. Young MD, Behjati S. SoupX removes ambient RNA contamination from droplet-based single-cell RNA sequencing data. Gigascience. 2020;9:giaa151.

42. Yang S, Corbett SE, Koga Y, Wang Z, Johnson WE, Yajima M, et al. Decontamination of ambient RNA in single-cell RNA-seq with DecontX. Genome Biology. 2020;21:57.

43. conos. 2023.

44. Jansky S, Sharma AK, Körber V, Quintero A, Toprak UH, Wecht EM, et al. Single-cell transcriptomic analyses provide insights into the developmental origins of neuroblastoma. Nat Genet. 2021;53:683–93.

45. Kildisiute G, Kholosy WM, Young MD, Roberts K, Elmentaite R, van Hooff SR, et al. Tumor to normal single-cell mRNA comparisons reveal a pan-neuroblastoma cancer cell. Sci Adv. 2021;7:eabd3311.

46. Han X, Zhou Z, Fei L, Sun H, Wang R, Chen Y, et al. Construction of a human cell landscape at single-cell level. Nature. 2020;581:303–9.

47. Torgersen S, Gjerdet NR, Erichsen ES, Bang G. Metal particles and tissue changes adjacent to miniplates. A retrieval study. Acta Odontol Scand. 1995;53:65–71.

48. Zhang Z, Zamojski M, Smith GR, Willis TL, Yianni V, Mendelev N, et al. Single nucleus transcriptome and chromatin accessibility of postmortem human pituitaries reveal diverse stem cell regulatory mechanisms. Cell Rep. 2022;38:110467.

49. La Manno G, Soldatov R, Zeisel A, Braun E, Hochgerner H, Petukhov V, et al. RNA velocity of single cells. Nature. 2018;560:494–8.

50. Wu T, Hu E, Xu S, Chen M, Guo P, Dai Z, et al. clusterProfiler 4.0: A universal enrichment tool for interpreting omics data. Innovation (Camb). 2021;2:100141.

51. Kiraz Y, Adan A, Kartal Yandim M, Baran Y. Major apoptotic mechanisms and genes involved in apoptosis. Tumour Biol. 2016;37:8471–86.

52. Nikoletopoulou V, Markaki M, Palikaras K, Tavernarakis N. Crosstalk between apoptosis, necrosis and autophagy. Biochim Biophys Acta. 2013;1833:3448–59.

53. Kanehisa M, Furumichi M, Sato Y, Ishiguro-Watanabe M, Tanabe M. KEGG: integrating viruses and cellular organisms. Nucleic Acids Res. 2021;49:D545–51.

54. Kanehisa M. Toward understanding the origin and evolution of cellular organisms. Protein Sci. 2019;28:1947–51.

55. Yer EN, Baloglu MC, Ayan S. Identification and expression profiling of all Hsp family member genes under salinity stress in different poplar clones. Gene. 2018;678:324–36.

56. Kampinga HH, Hageman J, Vos MJ, Kubota H, Tanguay RM, Bruford EA, et al. Guidelines for the nomenclature of the human heat shock proteins. Cell Stress Chaperones. 2009;14:105–11.

