## Additional file 1 for "Comparative framework and adaptation of ACME HS approach to single cell isolation from fresh-frozen endocrine tissues"

**Figure S1: RNA integrity and quantification.**

**a.** Correlative analysis of RIN values and the % area of the two ribosomal bands compared to the total.The linear correlation is indicated on the graph. **b.** Tables containing the individual values extracted from the RNA samples displayed in Figure 1 B, C, D and RNA cell samples of adrenocortical tumor, adrenal medullary tumor, thyroid carcinoma and PitNET.

**a**

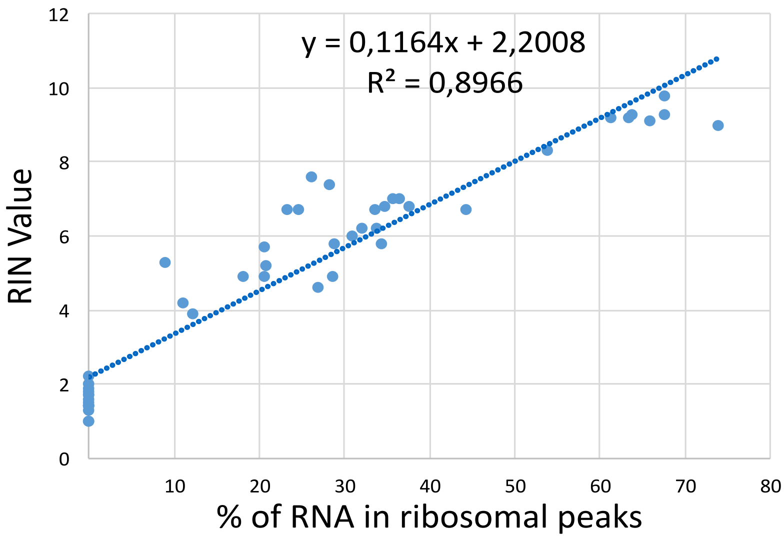


**b**

| Fig. 1B. RIN values of ACME HS-dissociated and cryopreserved cells obtained from adrenocortical tumor | | | |
|---|---|---|---|
| intervals | RIN | RNA conc (ng/ul) | % ribosomal RNA |
| 0 days | 7,6 | 53,8 | 26,2 |
| 1 day | 7,4 | 22,6 | 28,3 |
| 3 days | 7 | 14,2 | 35,7 |
| 7days | 6,7 | 25,6 | 24,6 |
| 14 days | 6,7 | 26,2 | 23,3 |
| 28 days | 5,7 | 11,5 | 20,5 |
| 6 mounths | 5,9 | 8,4 | 36,4 |

| Fig. 1B. RIN values of control adrenocortical tumor sample | | | |
|---|---|---|---|
| intervals | RIN | RNA conc (ng/ul) | % ribosomal RNA |
| 0 days | 9,8 | 53,8 | 67,5 |

| Fig. 1C. RIN values of enzyme-dissociated and methanol-fixed cells obtained from adrenocortical tumor | | | |
|---|---|---|---|
| intervals | RIN | conc (ng/ul) | % ribosomal RNA |
| 0 days | 8,3 | 4,5 | 53,8 |
| 1 day | 3,9 | 4,23 | 12,1 |
| 3 days | 2,2 | 4,23 | 0 |
| 7days | 1,7 | 24,5 | 0 |
| 14 days | 1,4 | 9,42 | 0 |
| 28 days | 1,5 | 7,05 | 0 |

| Fig. 1D. RIN values of fresh-frozen adrenocortical tumor, om which ACME HS-dissociated cells (Fig. 1B) were obtained | | | |
|---|---|---|---|
| intervals | RIN | RNA conc (ng/ul) | % ribosomal RNA |
| 0 days | 9,3 | 22,9 | 63,8 |
| 1 day | 9,3 | 71,0 | 67,6 |
| 3 days | 9,2 | 27,7 | 63,4 |
| 7days | 9.0 | 21,5 | 73,9 |
| 14 days | 9,1 | 14,2 | 65,9 |
| 28 days | 9,2 | 20,0 | 61,3 |

| RIN values of ACME HS-dissociated and cryopreserved cells obtained from adrenal medullary tumor | | | |
|---|---|---|---|
| intervals | RIN | RNA conc (ng/ul) | % ribosomal RNA |
| 0 days | 6,7 | 13,4 | 44,3 |
| 1 day | 5,8 | 31 | 34,4 |
| 3 days | 6,0 | 13,7 | 30,8 |
| 7days | 6,2 | 15,9 | 33,7 |
| 14 days | 6,2 | 13,9 | 32,1 |
| 28 days | 5,8 | 15,5 | 28,7 |

| RIN values of ACME HS-dissociated and cryopreserved cells obtained from thyroid carcinoma | | | |
|---|---|---|---|
| intervals | RIN | RNA conc (ng/ul) | % ribosomal RNA |
| 0 days | 6,7 | 53,2 | 33,5 |
| 1 day | 5,2 | 87 | 20,8 |
| 3 days | 4,9 | 39,6 | 20,6 |
| 7days | 4,9 | 80,2 | 28,6 |
| 14 days | 4,9 | 128 | 18,1 |
| 28 days | 4,6 | 130 | 26,9 |

| RIN values of ACME HS-dissociated and cryopreserved cells obtained from PitNET | | | |
|---|---|---|---|
| intervals | RIN | conc (ng/ul) | % ribosomal RNA |
| 0 days | 7 | 12,1 | 36,5 |
| 1 day | 6,8 | 65 | 34,7 |

| RIN values of enzyme-dissociated and methanol-fixed cells obtained from adrenal medullary tumor | | | |
|---|---|---|---|
| intervals | RIN | RNA conc (ng/ul) | % ribosomal RNA |
| 0 days | 5,3 | 4,43 | 9 |
| 1 day | 2 | 2,3 | 0 |
| 3 days | 1,8 | 3,24 | 0 |
| 7days | 1,9 | 11 | 0 |
| 14 days | 1,3 | 8,47 | 0 |
| 28 days | 1,4 | 1,95 | 0 |

| RIN values of enzyme-dissociated and methanol-fixed cells obtained from thyroid carcinoma | | | |
|---|---|---|---|
| intervals | RIN | RNA conc (ng/ul) | % ribosomal RNA |
| 0 days | 4,2 | 12,2 | 11 |
| 1 day | 1,6 | 1,5 | 0 |
| 3 days | 1,7 | 2,1 | 0 |
| 7days | 1,4 | 1,87 | 0 |
| 14 days | 1,0 | 2,35 | 0 |
| 28 days | 1,0 | 1,96 | 0 |

| RIN values of enzyme-dissociated and methanol-fixed cells obtained from PitNET | | | |
|---|---|---|---|
| intervals | RIN | conc (ng/ul) | % ribosomal RNA |
| 0 days | 6,8 | 1,94 | 37,5 |
| 1 day | 1,4 | 2,05 | 0 |

**Figure S2: Comparison of storage and morphology of ACME HS and enzyme-dissociated cells.** **a.** Gel image of isolated total RNA from cryopreserved ACME HS-dissociated and methanol-fixed enzyme-dissociated adrenocortical cells after 6 months of freezing at -80°C. **b**. Bright field (BF) and confocal fluorescence microscopy images of ACME HS and enzyme-dissociated adrenocortical and thyroid follicular cells stained with anti-CYP11B1 (red), anti-TSHR (red) antibody, respectively, and Hoechst 33342. **c.** Cell viability (%) data obtained from adrenal medullary tumors, adrenocortical tumors, thyroid carcinomas, and PitNETs according to the type of enzyme used. Values are plotted for each experiment, and mean ± SEM is indicated. Statistical differences estimated by t-test: ns - not significant – $p > 0.05$

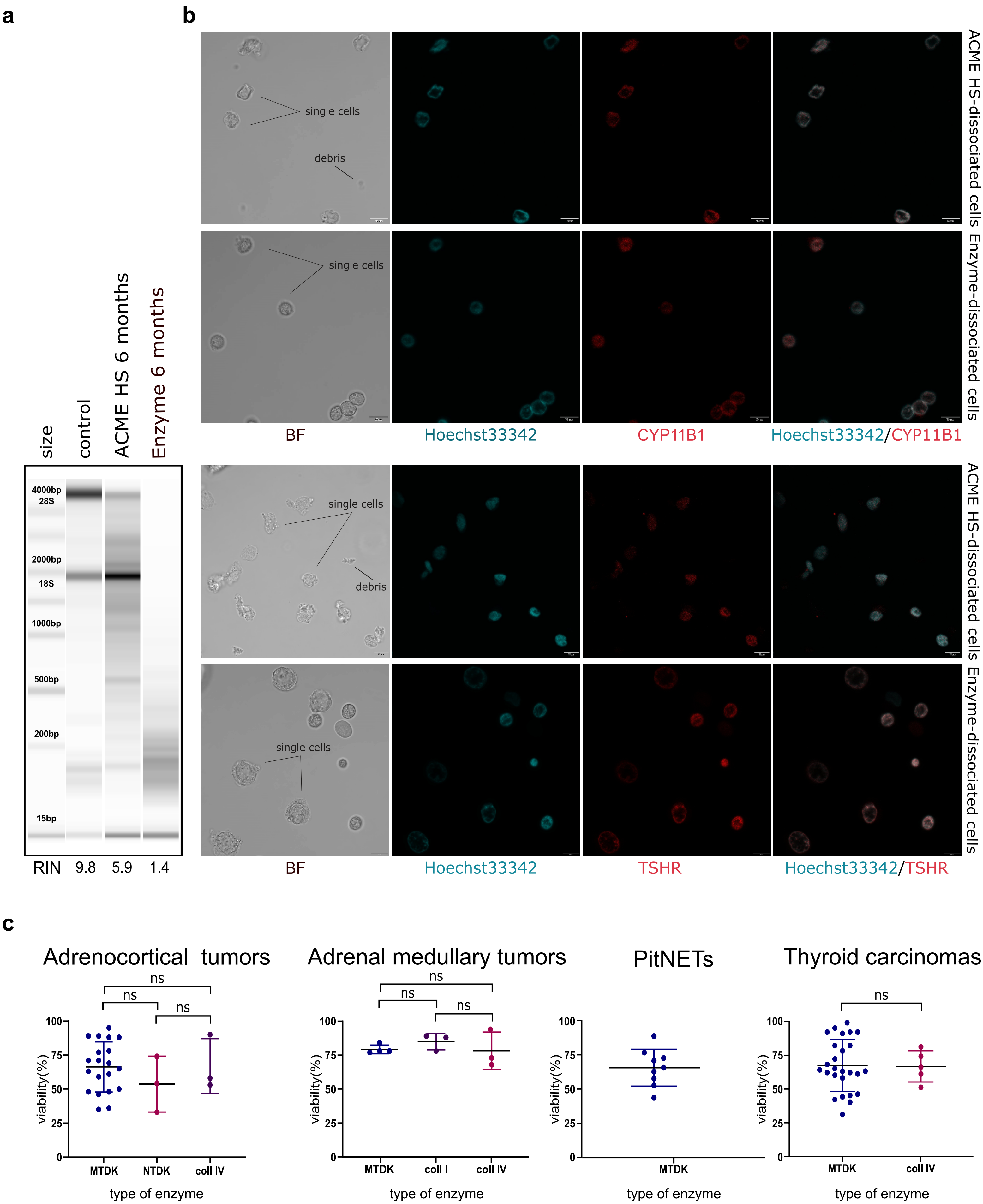
a
b
size
control
ACME HS 6 months
Enzyme 6 months
4000bp
28S
2000bp
18S
1000bp
500bp
200bp
15bp
RIN 9.8 5.9 1.4
single cells
debris
ACME HS-dissociated cells
Enzyme-dissociated cells
BF
Hoechst33342
CYP11B1
Hoechst33342/CYP11B1
TSHR
Hoechst33342/TSHR
c
Adrenocortical tumors
Adrenal medullary tumors
PitNETs
Thyroid carcinomas
ns
viability(%)
100
75
50
25
0
MTDK
NTDK
coll I
coll IV
type of enzyme

**Figure S3: Flow cytometry data of the different samples prepared through ACME HS method or enzymatic digestion**.

**a1-h1.** FSC-height/FSC-area dot plots are used to calculate cellular debris, single cells, and cellular aggregates (green events - for debris, red events - for singlets, and black events – for aggregates). **a2-h2.** FSC/SSC dot plots demonstrate the distribution of cells, their aggregates and cellular debris based on their light-scattering properties. **a3-h3**. PI-height/PI-area dot plots from all ungated events were used for additional location assessment for debris, singlets, and aggregates. **a4-h4**. PI-height/PI-area dot plots from singlets used for additional gating of single events (shown in blue) among nucleated cells and their aggregates (shown in red). **a5-h5.** DNA histograms from single events showing cell cycle distribution for all cells in the sample, with percentages of the cell cycle phases (G0/G1, S, G2/M) and mean fluorescence intensity for G0/G1 and G2/M phases inserted. The ACME HS method was performed in 3 replicates for each tissue.

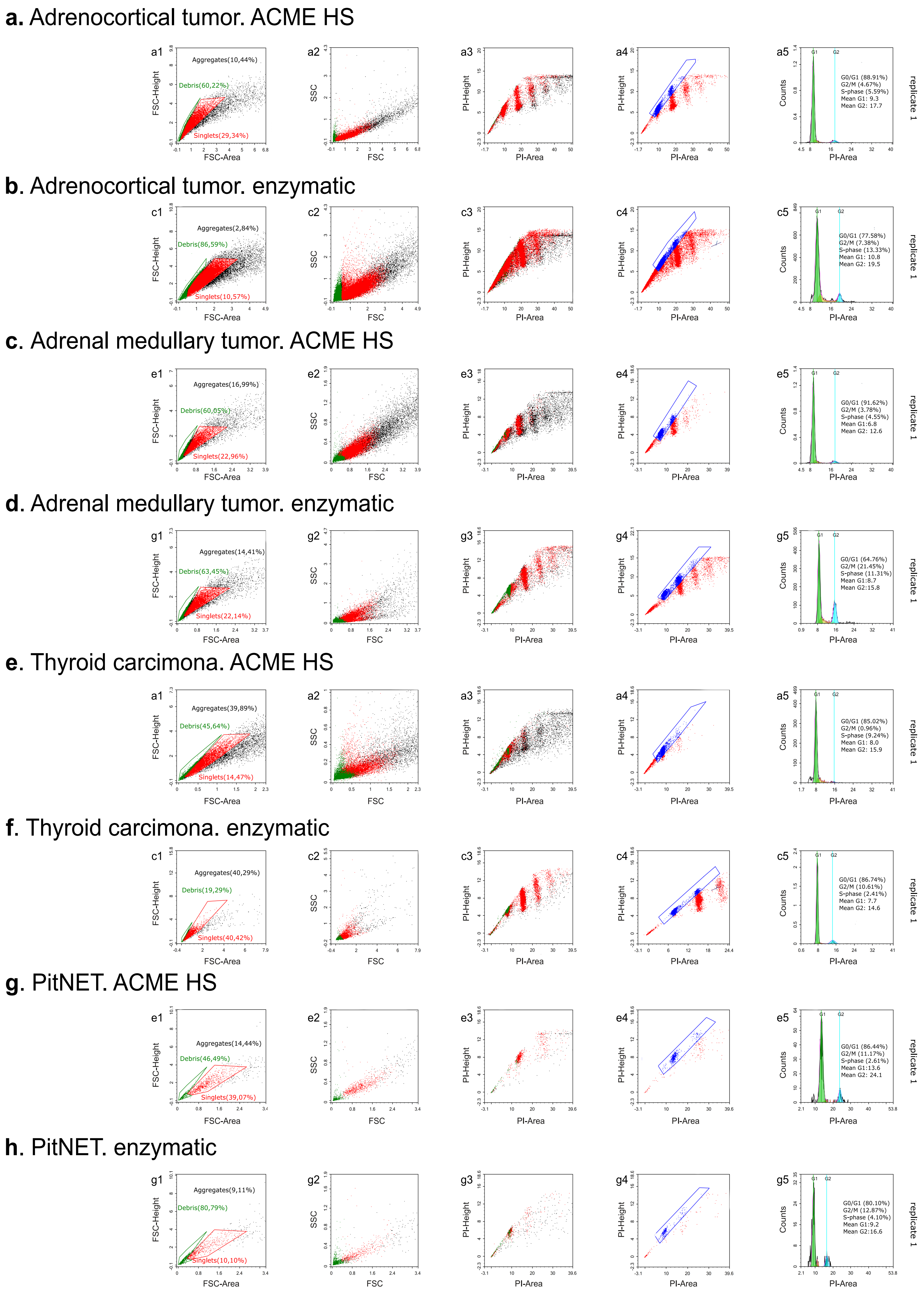
a. Adrenocortical tumor. ACME HS
a1
Aggregates(10,44%)
Debris(60,22%)
Singlets(29,34%)
FSC-Height
FSC-Area
a2
SSC
FSC
a3
PI-Height
PI-Area
a4
PI-Height
PI-Area
a5
G1
G2
G0/G1 (88.91%)
G2/M (4.67%)
S-phase (5.59%)
Mean G1: 9.3
Mean G2: 17.7
Counts
PI-Area
replicate 1
b. Adrenocortical tumor. enzymatic
c1
Aggregates(2,84%)
Debris(86,59%)
Singlets(10,57%)
FSC-Height
FSC-Area
c2
SSC
FSC
c3
PI-Height
PI-Area
c4
PI-Height
PI-Area
c5
G1
G2
G0/G1 (77.58%)
G2/M (7.38%)
S-phase (13.33%)
Mean G1: 10.8
Mean G2: 19.5
Counts
PI-Area
replicate 1
c. Adrenal medullary tumor. ACME HS
e1
Aggregates(16,99%)
Debris(60,05%)
Singlets(22,96%)
FSC-Height
FSC-Area
e2
SSC
FSC
e3
PI-Height
PI-Area
e4
PI-Height
PI-Area
e5
G1
G2
G0/G1 (91.62%)
G2/M (3.78%)
S-phase (4.55%)
Mean G1: 6.8
Mean G2: 12.6
Counts
PI-Area
replicate 1
d. Adrenal medullary tumor. enzymatic
g1
Aggregates(14,41%)
Debris(63,45%)
Singlets(22,14%)
FSC-Height
FSC-Area
g2
SSC
FSC
g3
PI-Height
PI-Area
g4
PI-Height
PI-Area
g5
G1
G2
G0/G1 (64.76%)
G2/M (21.45%)
S-phase (11.31%)
Mean G1:8.7
Mean G2:15.8
Counts
PI-Area
replicate 1
e. Thyroid carcimona. ACME HS
a1
Aggregates(39,89%)
Debris(45,64%)
Singlets(14,47%)
FSC-Height
FSC-Area
a2
SSC
FSC
a3
PI-Height
PI-Area
a4
PI-Height
PI-Area
a5
G1
G2
G0/G1 (85.02%)
G2/M (0.96%)
S-phase (9.24%)
Mean G1: 8.0
Mean G2: 15.9
Counts
PI-Area
replicate 1
f. Thyroid carcimona. enzymatic
c1
Aggregates(40,29%)
Debris(19,29%)
Singlets(40,42%)
FSC-Height
FSC-Area
c2
SSC
FSC
c3
PI-Height
PI-Area
c4
PI-Height
PI-Area
c5
G1
G2
G0/G1 (86.74%)
G2/M (10.61%)
S-phase (2.41%)
Mean G1: 7.7
Mean G2: 14.6
Counts
PI-Area
replicate 1
g. PitNET. ACME HS
e1
Aggregates(14,44%)
Debris(46,49%)
Singlets(39,07%)
FSC-Height
FSC-Area
e2
SSC
FSC
e3
PI-Height
PI-Area
e4
PI-Height
PI-Area
e5
G1
G2
G0/G1 (86.44%)
G2/M (11.17%)
S-phase (2.61%)
Mean G1:13.6
Mean G2: 24.1
Counts
PI-Area
replicate 1
h. PitNET. enzymatic
g1
Aggregates(9,11%)
Debris(80,79%)
Singlets(10,10%)
FSC-Height
FSC-Area
g2
SSC
FSC
g3
PI-Height
PI-Area
g4
PI-Height
PI-Area
g5
G1
G2
G0/G1 (80.10%)
G2/M (12.87%)
S-phase (4.10%)
Mean G1:9.2
Mean G2:16.6
Counts
PI-Area
replicate 1

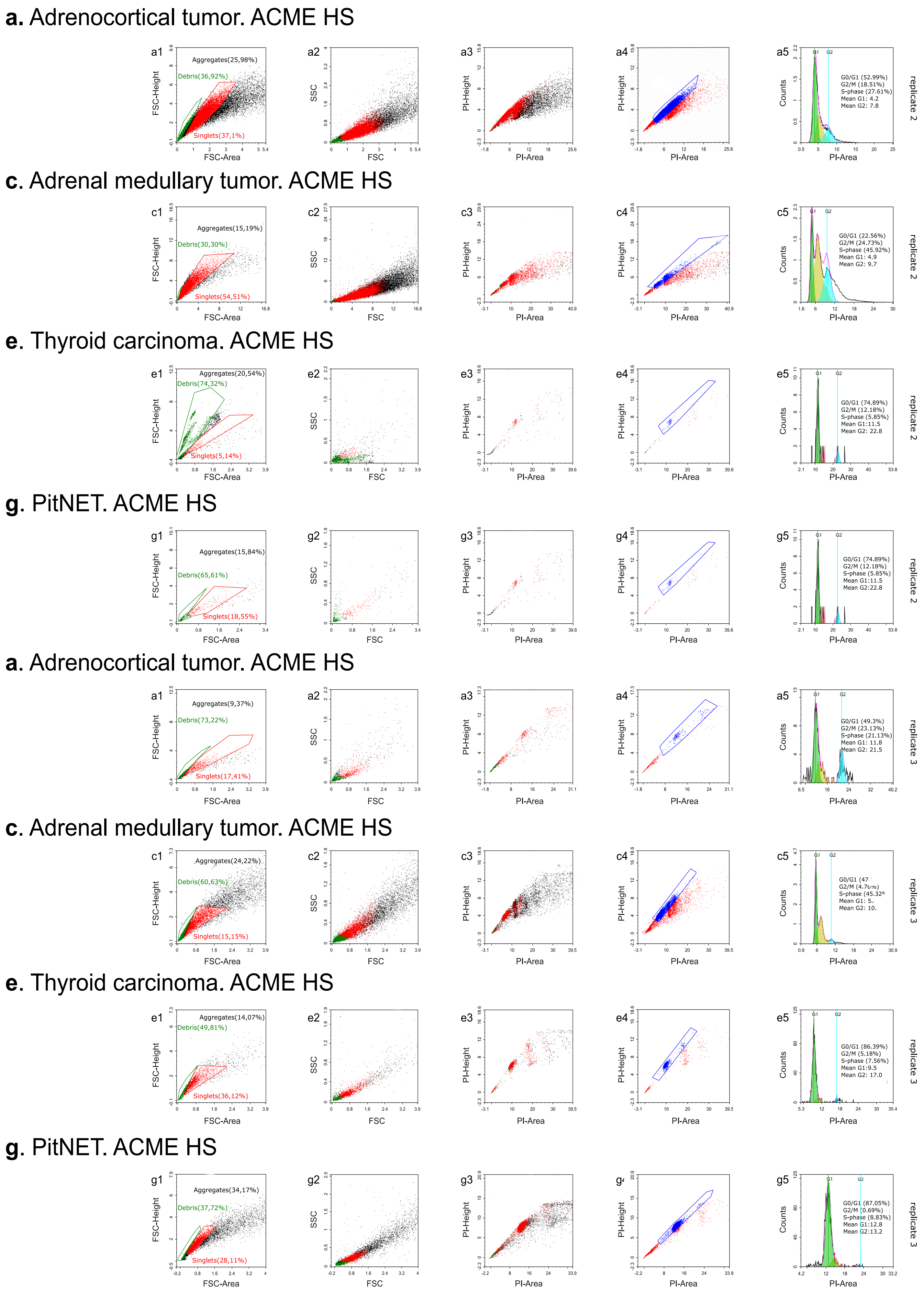

a. Adrenocortical tumor. ACME HS
Aggregates(25,98%)
Debris(36,92%)
Singlets(37,1%)
G0/G1 (52.99%)
G2/M (18.51%)
S-phase (27.61%)
Mean G1: 4.2
Mean G2: 7.8
replicate 2
c. Adrenal medullary tumor. ACME HS
Aggregates(15,19%)
Debris(30,30%)
Singlets(54,51%)
G0/G1 (22.56%)
G2/M (24.73%)
S-phase (45.92%)
Mean G1: 4.9
Mean G2: 9.7
replicate 2
e. Thyroid carcinoma. ACME HS
Aggregates(20,54%)
Debris(74,32%)
Singlets(5,14%)
G0/G1 (74.89%)
G2/M (12.18%)
S-phase (5.85%)
Mean G1:11.5
Mean G2: 22.8
replicate 2
g. PitNET. ACME HS
Aggregates(15,84%)
Debris(65,61%)
Singlets(18,55%)
G0/G1 (74.89%)
G2/M (12.18%)
S-phase (5.85%)
Mean G1:11.5
Mean G2:22.8
replicate 2
a. Adrenocortical tumor. ACME HS
Aggregates(9,37%)
Debris(73,22%)
Singlets(17,41%)
G0/G1 (49.3%)
G2/M (23.13%)
S-phase (21.13%)
Mean G1: 11.8
Mean G2: 21.5
replicate 3
c. Adrenal medullary tumor. ACME HS
Aggregates(24,22%)
Debris(60,63%)
Singlets(15,15%)
replicate 3
e. Thyroid carcinoma. ACME HS
Aggregates(14,07%)
Debris(49,81%)
Singlets(36,12%)
G0/G1 (86.39%)
G2/M (5.18%)
S-phase (7.56%)
Mean G1:9.5
Mean G2: 17.0
replicate 3
g. PitNET. ACME HS
Aggregates(34,17%)
Debris(37,72%)
Singlets(28,11%)
G0/G1 (87.05%)
G2/M (0.69%)
S-phase (8.83%)
Mean G1:12.8
Mean G2:13.2
replicate 3
FSC-Height
FSC-Area
SSC
FSC
PI-Height
PI-Area
Counts

**Figure S4: Microscopic images of endocrine tumor samples.**

**a.** Gross image of the adrenocortical tumor, showing foci of necrosis. **b.** A microscopic image of the adrenocortical tumor with foci of necrosis, x100, H&E. **c.** Gross image of the adrenal medullary tumor, showing foci of fibrosis. **d.** A microscopic image of the adrenal medullary tumor with foci of fibrosis, x100, H&E. E. Gross image of thyroid carcinoma. **f.** A microscopic image of the thyroid carcinoma with extensive amyloid deposition, x100, H&E. **g.** Gross image of PitNET. **h.** Microscopic image of the PitNET, x100, H&E. *Note*: the green dotted line – foci of necrosis; the blue dotted line – showing foci of fibrosis and amyloid deposition; the red dotted line – normal pituitary tissue.

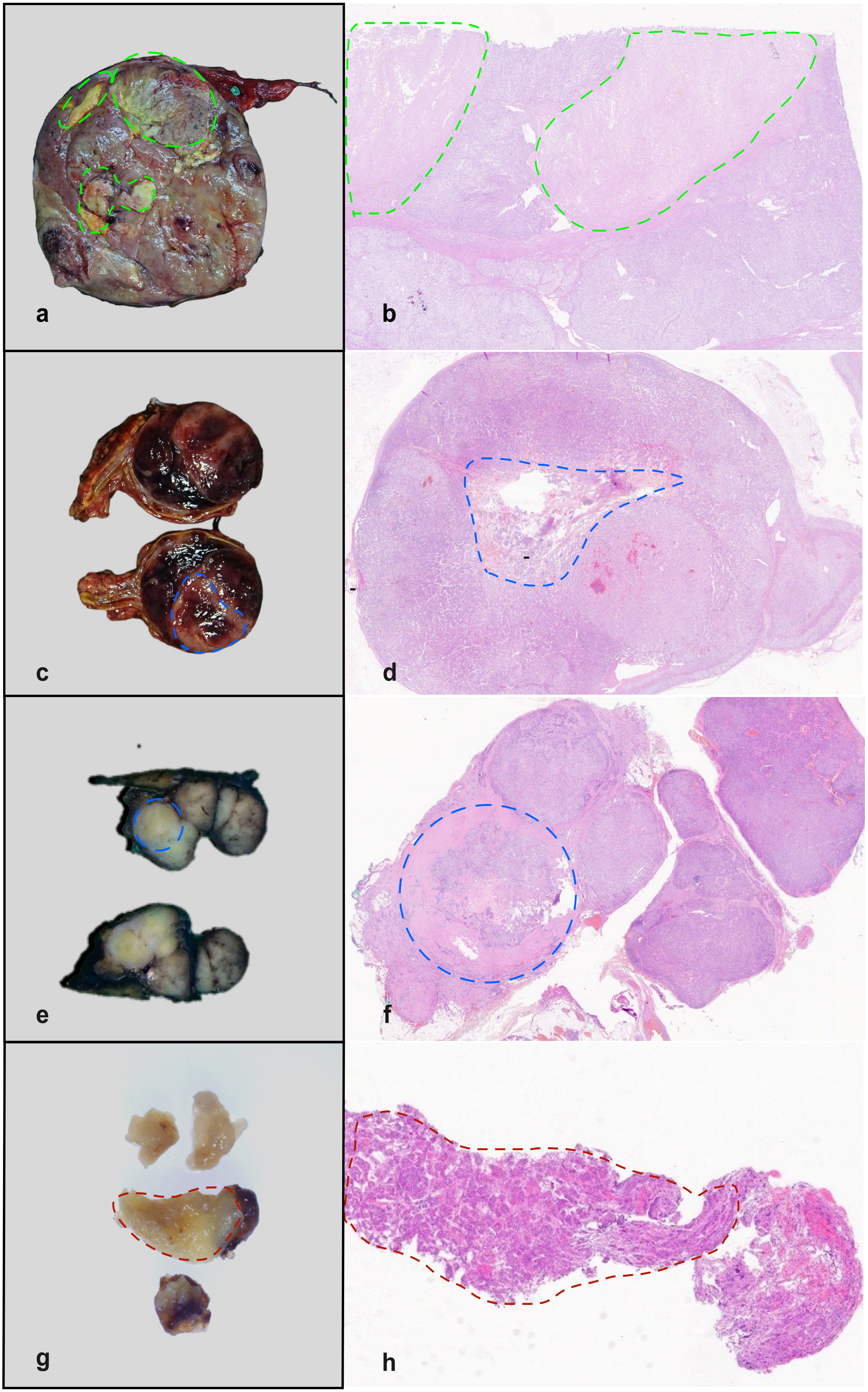
a
b
c
d
e
f
g
h

**Figure S5: Single-cell sample preparation methods comparison**.

**a**, **b**. Comparison of the basic sample features calculated by the Cellranger pipeline. **c.** Standard single-cell sample features comparisons (number of reads, expressed genes, doublets, etc.) for ACME HS, enzyme and nuclei isolation methods. **d.** Standard single-cell sample features comparisons of all four tissues (adrenocortical tumors, adrenal medullary tumors, thyroid carcinomas and PitNETs) for each dissociation method. Wilcoxon rank sum used for statistics calculation: **** ($0.0001 < p < 0.001$), *** ($p < 0.001$), ** ($0.001 < p < 0.01$), * ($0.01 < p < 0.05$), ns - not significant – $p > 0.05$.

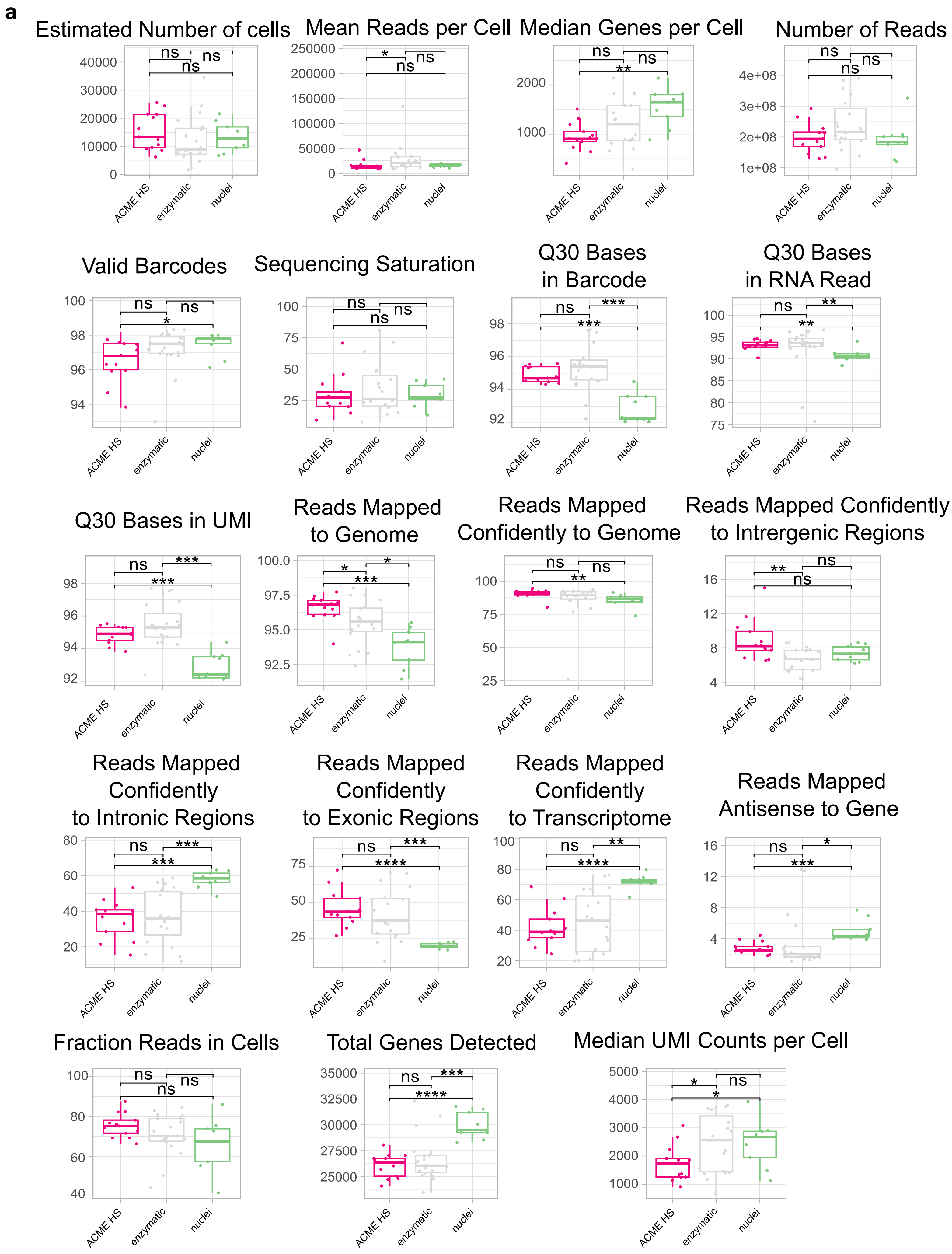
a
Estimated Number of cells
0
10000
20000
30000
40000
Mean Reads per Cell
0
50000
100000
150000
200000
250000
Median Genes per Cell
1000
2000
Number of Reads
1e+08
2e+08
3e+08
4e+08
Valid Barcodes
94
96
98
100
Sequencing Saturation
25
50
75
100
Q30 Bases in Barcode
92
94
96
98
Q30 Bases in RNA Read
75
80
85
90
95
100
Q30 Bases in UMI
92
94
96
98
Reads Mapped to Genome
92.5
95.0
97.5
100.0
Reads Mapped Confidently to Genome
25
50
75
100
Reads Mapped Confidently to Intrergenic Regions
4
8
12
16
Reads Mapped Confidently to Intronic Regions
20
40
60
80
Reads Mapped Confidently to Exonic Regions
25
50
75
Reads Mapped Confidently to Transcriptome
20
40
60
80
100
Reads Mapped Antisense to Gene
4
8
12
16
Fraction Reads in Cells
40
60
80
100
Total Genes Detected
25000
27500
30000
32500
35000
Median UMI Counts per Cell
1000
2000
3000
4000
5000
ACME HS
enzymatic
nuclei
ns
*
**
***
****

**b**

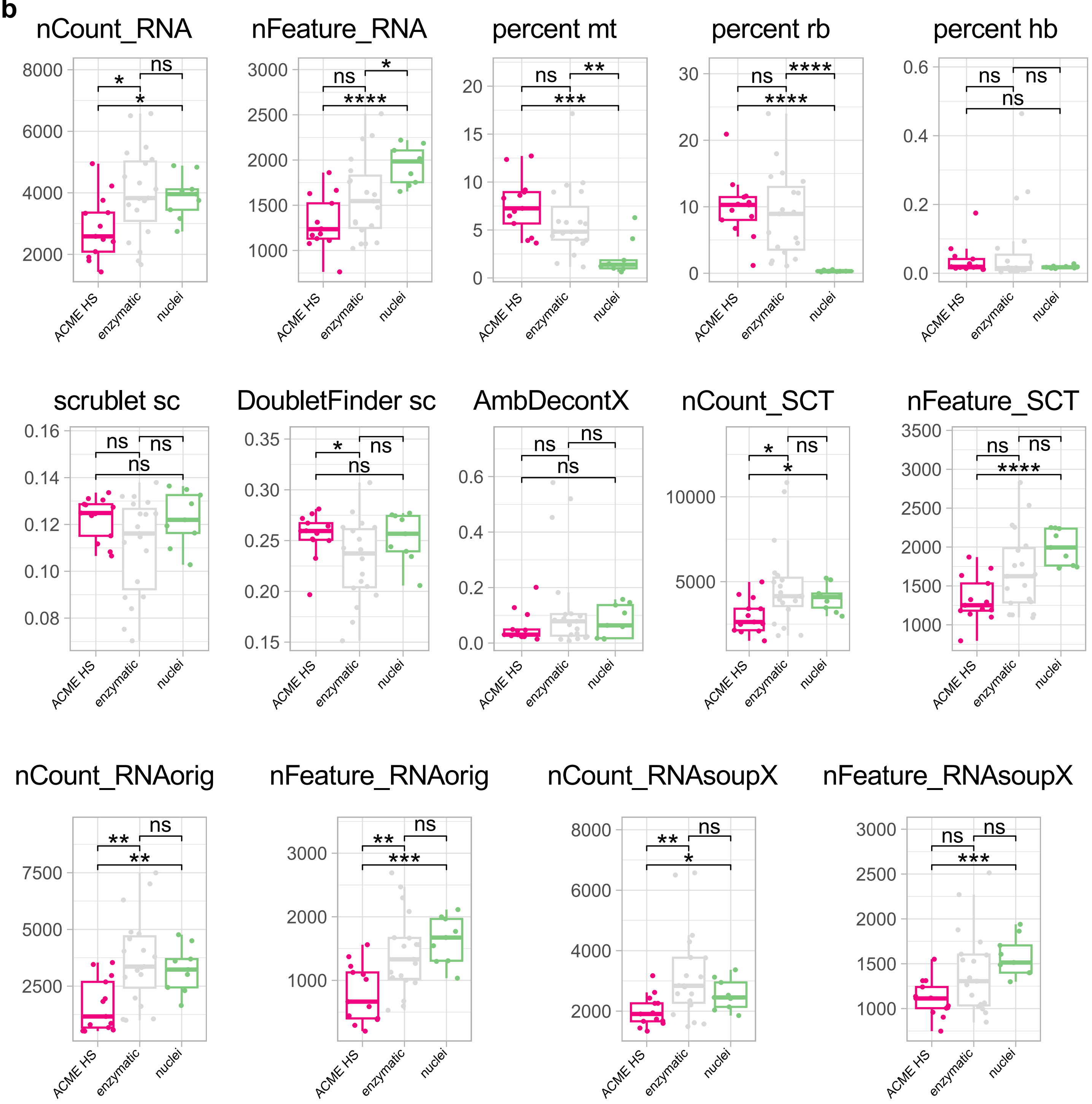
nCount_RNA
nFeature_RNA
percent mt
percent rb
percent hb
scrublet sc
DoubletFinder sc
AmbDecontX
nCount_SCT
nFeature_SCT
nCount_RNAorig
nFeature_RNAorig
nCount_RNAsoupX
nFeature_RNAsoupX
ACME HS
enzymatic
nuclei
ns
*
**
***
****

c

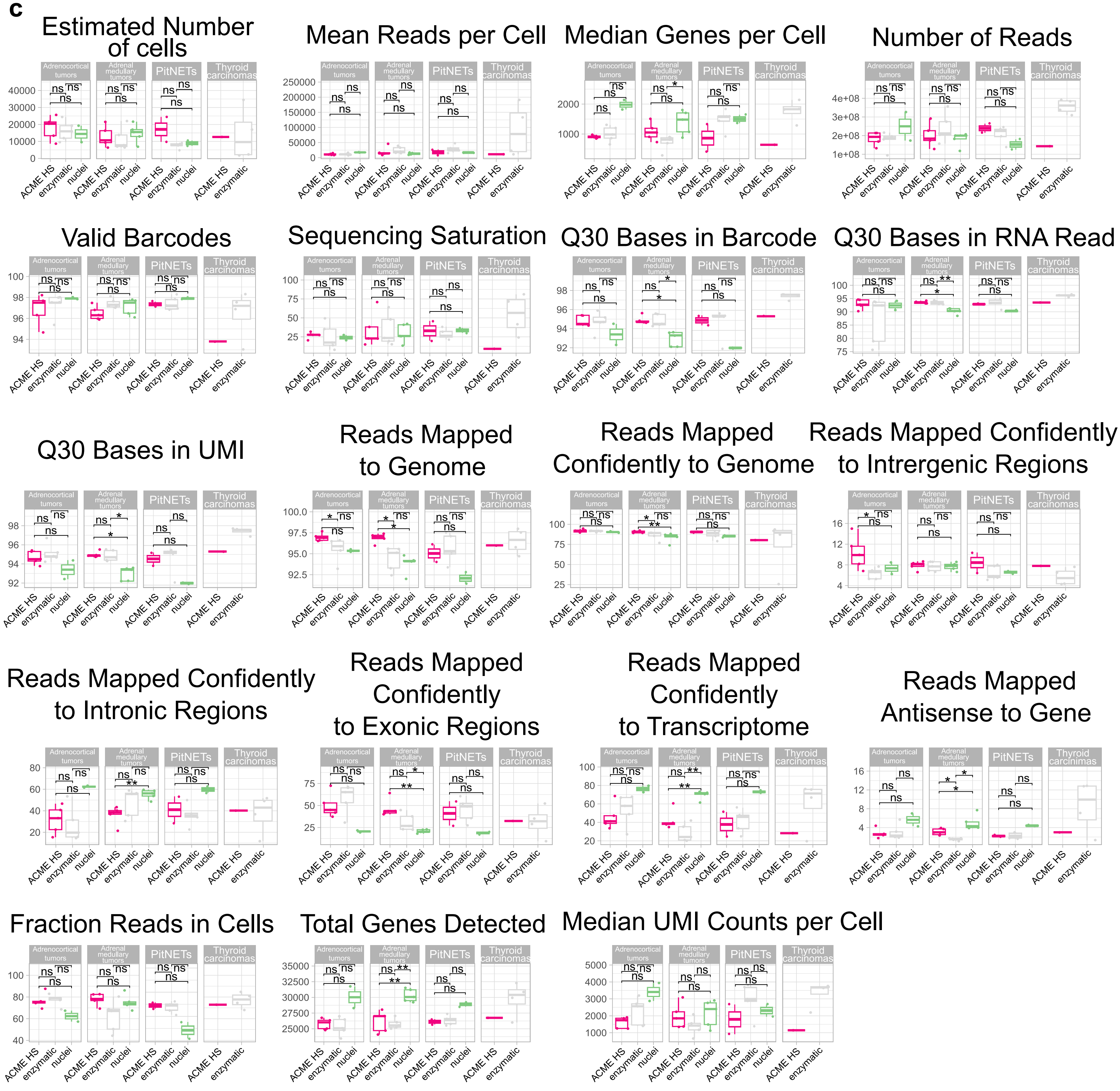
Estimated Number of cells
Mean Reads per Cell
Median Genes per Cell
Number of Reads
Valid Barcodes
Sequencing Saturation
Q30 Bases in Barcode
Q30 Bases in RNA Read
Q30 Bases in UMI
Reads Mapped to Genome
Reads Mapped Confidently to Genome
Reads Mapped Confidently to Intrergenic Regions
Reads Mapped Confidently to Intronic Regions
Reads Mapped Confidently to Exonic Regions
Reads Mapped Confidently to Transcriptome
Reads Mapped Antisense to Gene
Fraction Reads in Cells
Total Genes Detected
Median UMI Counts per Cell
Adrenocortical tumors
Adrenal medullary tumors
PitNETs
Thyroid carcinomas
Thyroid carcinimas
ACME HS
enzymatic
nuclei
ns
*
**

d

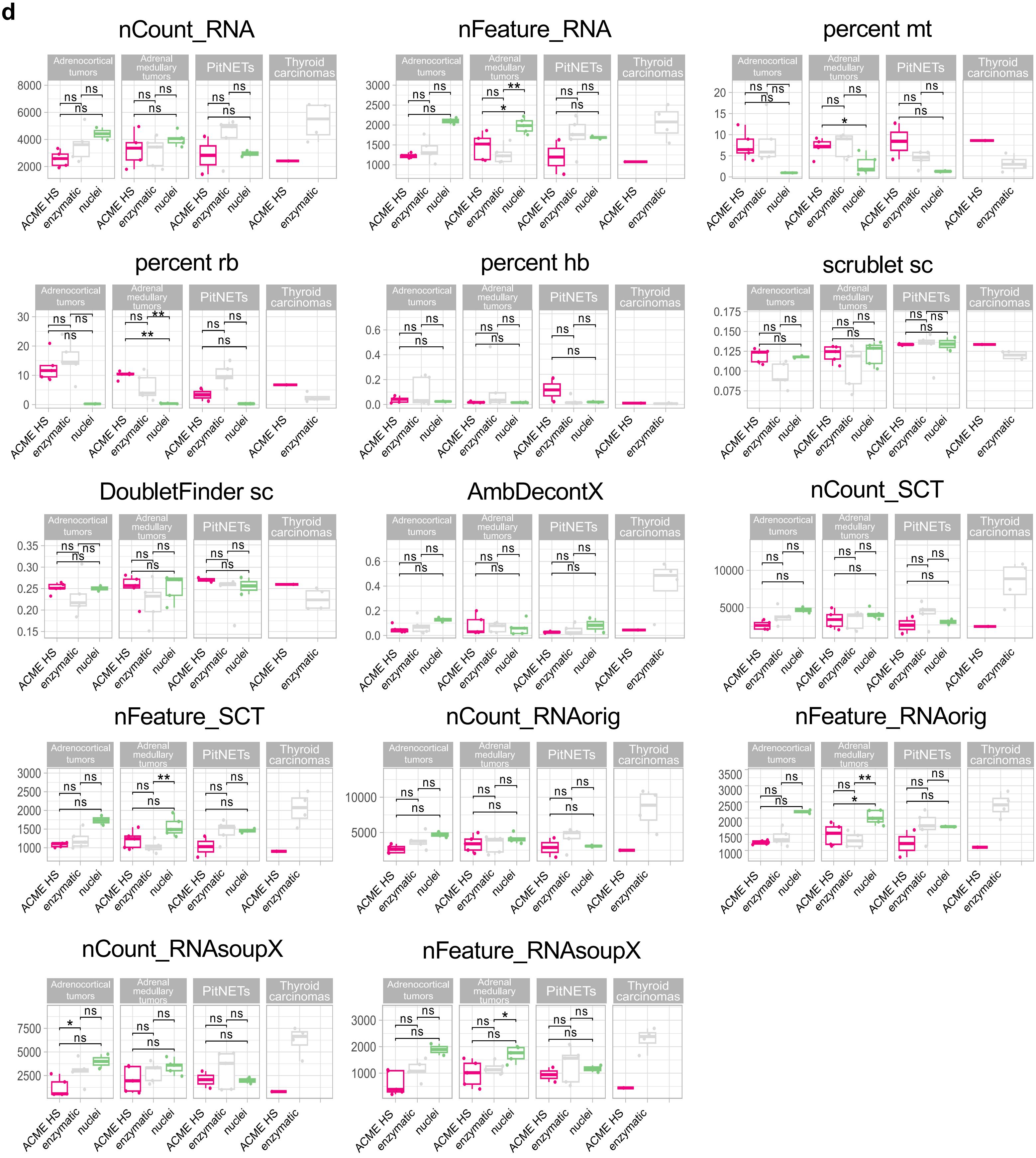
nCount_RNA
nFeature_RNA
percent mt
percent rb
percent hb
scrublet sc
DoubletFinder sc
AmbDecontX
nCount_SCT
nFeature_SCT
nCount_RNAorig
nFeature_RNAorig
nCount_RNAsoupX
nFeature_RNAsoupX
Adrenocortical tumors
Adrenal medullary tumors
PitNETs
Thyroid carcinomas
ACME HS
enzymatic
nuclei
ns
*
**

**Figure S6: Heterogeneity of major cell types in adrenocortical tumor and adrenal medullary tumor samples**. **a.** Visualization of the major cell subpopulations and states. **b.** Fractions of defined subpopulations and states identified by different methods. Adrenocortical and chromaffin cells were segregated by cell clustering applied on integrated data sets. Cells segregated into small clusters (<100 cells) were combined into separate minor groups and excluded from the analysis. The diagram does not indicate the number of cells representing less than 5% of the total number.

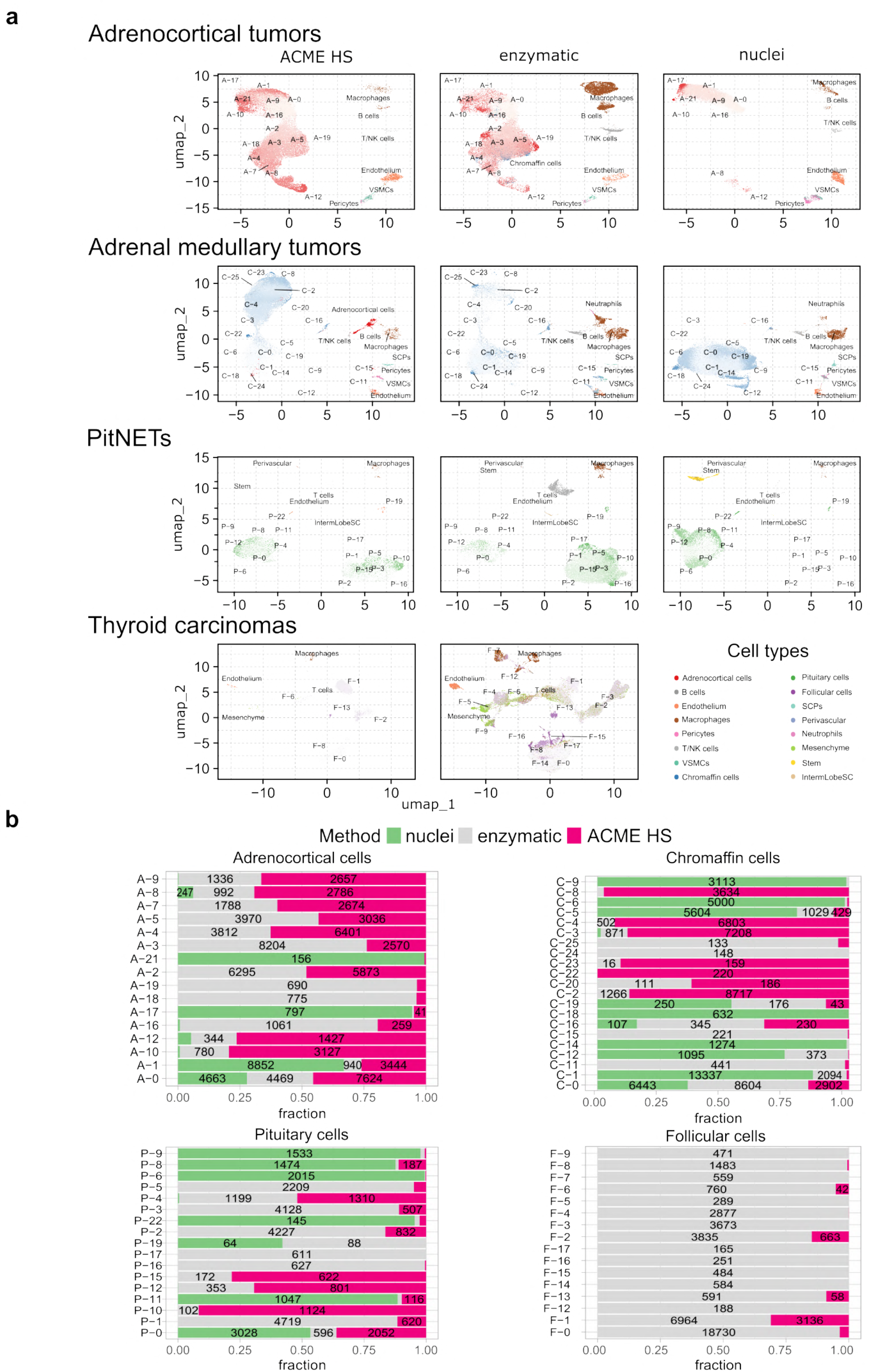
a
Adrenocortical tumors
ACME HS
enzymatic
nuclei
Adrenal medullary tumors
PitNETs
Thyroid carcinomas
umap_2
umap_1
Cell types
Adrenocortical cells
B cells
Endothelium
Macrophages
Pericytes
T/NK cells
VSMCs
Chromaffin cells
Pituitary cells
Follicular cells
SCPs
Perivascular
Neutrophils
Mesenchyme
Stem
IntermLobeSC
b
Method nuclei enzymatic ACME HS
Adrenocortical cells
Chromaffin cells
Pituitary cells
Follicular cells
fraction

**Figure S7: Functional characterization of cell clusters specific for ACME HS and enzymatic sample preparation methods.** Differentially expressed genes (up-regulated in selected clusters compared to all the rest cells, logFC > 0.25 and FDR < 0.05) were calculated for each cluster within major cell types **(Supplementary Fig. 6)** and analyzed by using wiki pathways as a reference database. **a**. Adrenocortical tumors. **b**. Adrenal medullary tumors. **c**. Alignment of the cells derived by ACME HS with enzymatic and nuclei data sets. ACME HS cells colored red, blue, green and purple for adrenocortical, chromaffin, thyroid follicular cells, and pituicytes, respectively. Cells derived by enzymatic and nuclei methods colored gray.

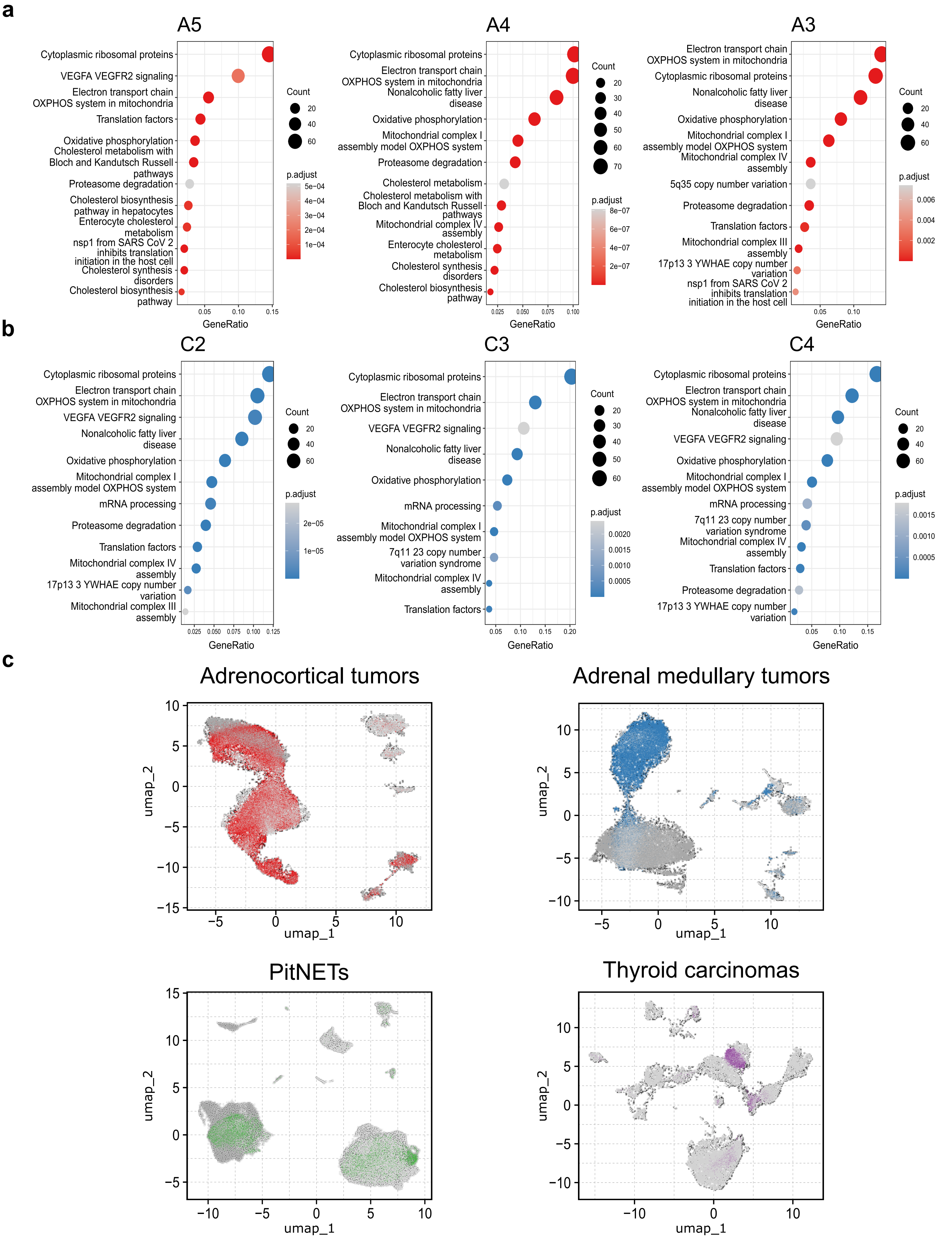
a
A5
Cytoplasmic ribosomal proteins
VEGFA VEGFR2 signaling
Electron transport chain OXPHOS system in mitochondria
Translation factors
Oxidative phosphorylation
Cholesterol metabolism with Bloch and Kandutsch Russell pathways
Proteasome degradation
Cholesterol biosynthesis pathway in hepatocytes
Enterocyte cholesterol metabolism
nsp1 from SARS CoV 2 inhibits translation initiation in the host cell
Cholesterol synthesis disorders
Cholesterol biosynthesis pathway
GeneRatio
Count
p.adjust
A4
Cytoplasmic ribosomal proteins
Electron transport chain OXPHOS system in mitochondria
Nonalcoholic fatty liver disease
Oxidative phosphorylation
Mitochondrial complex I assembly model OXPHOS system
Proteasome degradation
Cholesterol metabolism
Cholesterol metabolism with Bloch and Kandutsch Russell pathways
Mitochondrial complex IV assembly
Enterocyte cholesterol metabolism
Cholesterol synthesis disorders
Cholesterol biosynthesis pathway
GeneRatio
Count
p.adjust
A3
Electron transport chain OXPHOS system in mitochondria
Cytoplasmic ribosomal proteins
Nonalcoholic fatty liver disease
Oxidative phosphorylation
Mitochondrial complex I assembly model OXPHOS system
Mitochondrial complex IV assembly
5q35 copy number variation
Proteasome degradation
Translation factors
Mitochondrial complex III assembly
17p13 3 YWHAE copy number variation
nsp1 from SARS CoV 2 inhibits translation initiation in the host cell
GeneRatio
Count
p.adjust
b
C2
Cytoplasmic ribosomal proteins
Electron transport chain OXPHOS system in mitochondria
VEGFA VEGFR2 signaling
Nonalcoholic fatty liver disease
Oxidative phosphorylation
Mitochondrial complex I assembly model OXPHOS system
mRNA processing
Proteasome degradation
Translation factors
Mitochondrial complex IV assembly
17p13 3 YWHAE copy number variation
Mitochondrial complex III assembly
GeneRatio
Count
p.adjust
C3
Cytoplasmic ribosomal proteins
Electron transport chain OXPHOS system in mitochondria
VEGFA VEGFR2 signaling
Nonalcoholic fatty liver disease
Oxidative phosphorylation
mRNA processing
Mitochondrial complex I assembly model OXPHOS system
7q11 23 copy number variation syndrome
Mitochondrial complex IV assembly
Translation factors
GeneRatio
Count
p.adjust
C4
Cytoplasmic ribosomal proteins
Electron transport chain OXPHOS system in mitochondria
Nonalcoholic fatty liver disease
VEGFA VEGFR2 signaling
Oxidative phosphorylation
Mitochondrial complex I assembly model OXPHOS system
mRNA processing
7q11 23 copy number variation syndrome
Mitochondrial complex IV assembly
Translation factors
Proteasome degradation
17p13 3 YWHAE copy number variation
GeneRatio
Count
p.adjust
c
Adrenocortical tumors
Adrenal medullary tumors
PitNETs
Thyroid carcinomas
umap_1
umap_2

**Figure S8: Top gene examples contributing to the differences between tested methods.**
**a.** Gene expression changes for selected gene examples are visualized on UMAP namely, *CYP17A1* and *CYB5A* for adrenocortical tumor; **b –** *SYP*, *DBH* and *PNMT* for adrenal medullary tumor; **c –** *GH1* and *POU1F1* for PitNET; **d –** *TSHR* for thyroid carcinoma samples. **c.** Top differentially expressed genes (n=20) shown on the dotplots. **e.** For differential expression analysis ACME HS was compared against enzymatic and nuclei methods.

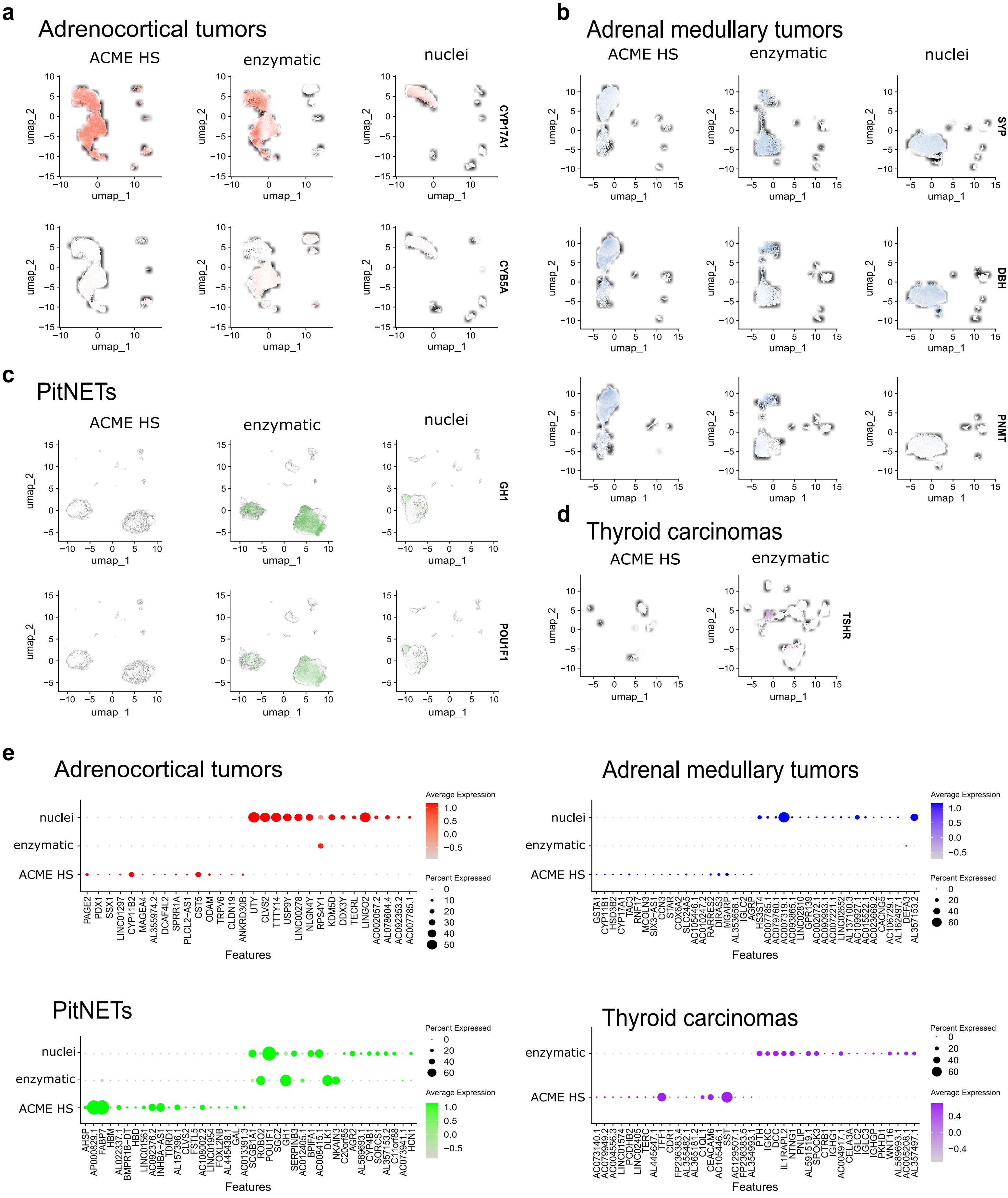

a
Adrenocortical tumors
ACME HS
enzymatic
nuclei
CYP17A1
CYB5A
umap_1
umap_2
b
Adrenal medullary tumors
SYP
DBH
PNMT
c
PitNETs
GH1
POU1F1
d
Thyroid carcinomas
TSHR
e
Adrenocortical tumors
Average Expression
Percent Expressed
Features
PAGE2
PDX1
SSX1
LINC01297
CYP11B2
MAGEA4
AL355974.2
DCAF4L2
SPRR1A
PLCL2-AS1
CST5
ODAM
TRPV6
CLDN19
ANKRD30B
UTY
CLVS2
TTTY14
USP9Y
LINC00278
NLGN4Y
RPS4Y1
KDM5D
DDX3Y
TECRL
LINGO2
AC002057.2
AL078604.4
AC092353.2
AC007785.1
Adrenal medullary tumors
GSTA1
CYP11B1
HSD3B2
CYP17A1
TAC3
RNF17
MCOLN3
SIX3-AS1
CCN3
STAR
COX6A2
SLC24A5
AC105446.1
AC010247.2
RARRES2
DIRAS3
MGARP
AL353658.1
IGLC2
AGRP
HS3ST4
AC007785.1
AC079760.1
AC007319.1
AC093865.1
LINC02810
GPR139
AC002072.2
AC090993.1
AC007221.1
LINC02652
AL137100.3
AC109927.1
AC015522.1
AC023669.2
CACNG5
AC106729.1
AL162497.1
DEFA3
AL357153.2
PitNETs
AHSP
AP000829.1
FABP7
HBM
AL022337.1
BMPR1B-DT
HBD
LINC01561
AC092376.2
INHBA-AS1
TDRD1
AL157396.1
CLVS2
FSTL5
AC108002.2
LINC01954
FOXL2NB
AL445438.1
GAL
AC013391.3
SCGB1A1
ROBO2
POU1F1
SGCZ
GH1
SERPINB3
AC012405.1
BPIFA1
AC008415.1
DLK1
NKAIN3
C20orf85
AGR2
AL589693.1
CYP4B1
SORCS1
AL357153.2
C11orf88
AC073941.1
HCN1
Thyroid carcinomas
AC073140.1
AC079949.2
AC004556.3
LINC01574
PCDHB2
LINC02405
TERT
AL445647.1
TFF1
CDR1
FP236383.4
AL355482.1
AL365181.2
C1QL1
CEACAM6
AC105446.1
SST
AC129507.1
FP236383.5
AL354993.1
PTH
IGKC
DCC
IL1RAPL2
NTNG1
PNLIP
AL591501.1
SPOCK3
CTRB1
IGHG1
AC004917.1
CELA3A
IGLC2
IGLC3
IGHGP
PKHD1
WNT16
AL589693.1
AC005208.1
AL357497.1

**Figure S9: Distribution of the enrichment scores of stress-related gene signatures across sample preparation methods.** The boxplots included inside the violin plots summarize the data distribution. Upper and lower sides of the box represent the 1st and 3rd quartiles. The line in the middle corresponds to the median. Lines extend no further than 1.5 the interquartile range. In all cases, statistical significance was tested using a one-tailed Wilcoxon rank-sum test: *** ($p < 0.001$), * ($0.01 < p < 0.05$), ns - not significant – $p > 0.05$. **a, b, c, d.** The stress signatures for adrenocortical tumor (n=12), adrenal medullary tumor (n=15), thyroid carcinoma (n=8) and PitNET (n=9) samples, respectively.

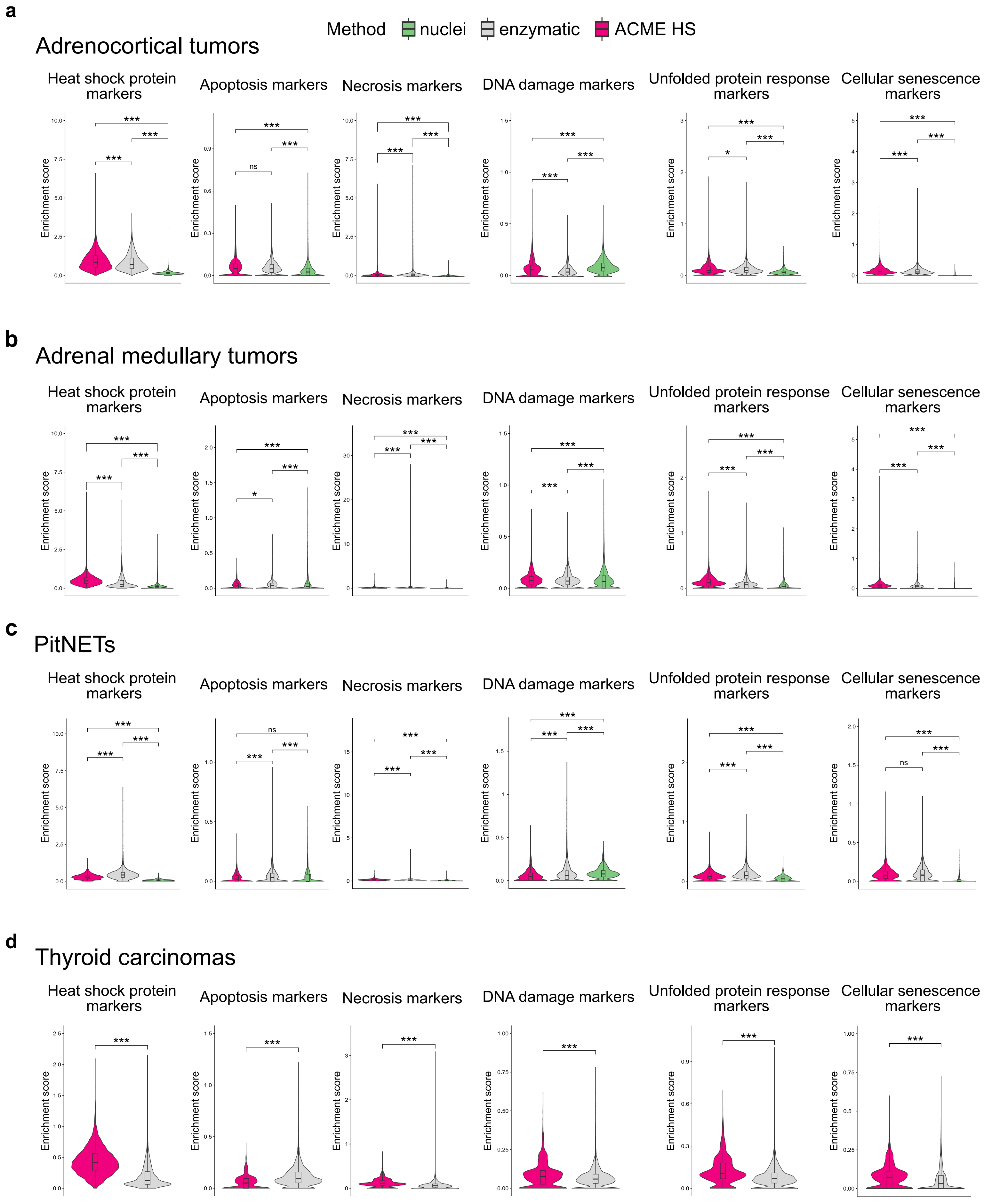
a
Method
nuclei
enzymatic
ACME HS
Adrenocortical tumors
Heat shock protein markers
Apoptosis markers
Necrosis markers
DNA damage markers
Unfolded protein response markers
Cellular senescence markers
Enrichment score
b
Adrenal medullary tumors
Heat shock protein markers
Apoptosis markers
Necrosis markers
DNA damage markers
Unfolded protein response markers
Cellular senescence markers
Enrichment score
c
PitNETs
Heat shock protein markers
Apoptosis markers
Necrosis markers
DNA damage markers
Unfolded protein response markers
Cellular senescence markers
Enrichment score
d
Thyroid carcinomas
Heat shock protein markers
Apoptosis markers
Necrosis markers
DNA damage markers
Unfolded protein response markers
Cellular senescence markers
Enrichment score

**Figure S10: Velocity and cell cycle estimations for PitNET datasets.**Velocity and cell cycle estimation for PitNETs datasets. Velocity was performed for individual samples (n=1) for each method, cell cycle estimation for PitNETs (n=9). Examples of differentially expressed (DE) genes associated with cell cycle control are shown on the individual embeddings.

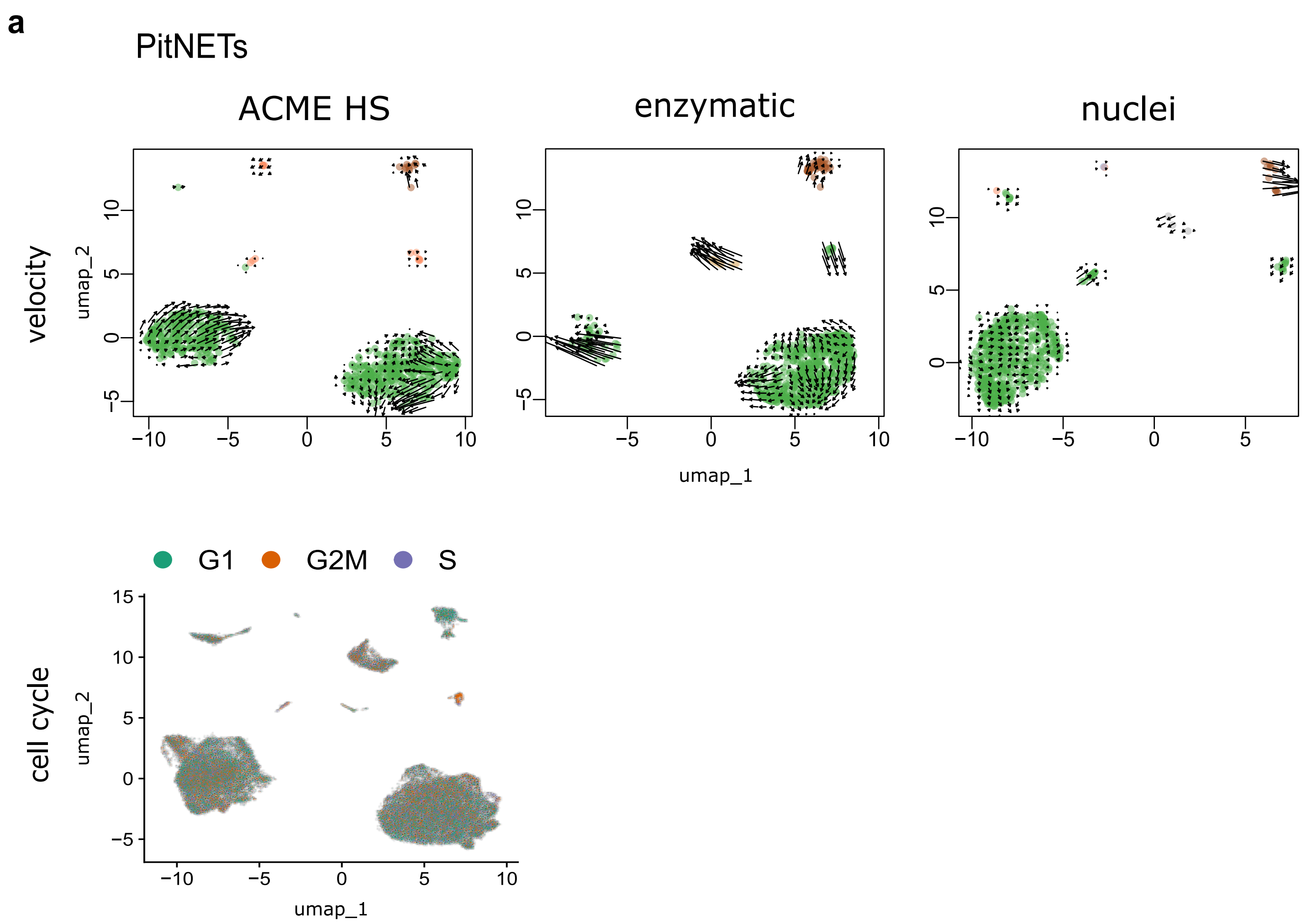

a
PitNETs
ACME HS
enzymatic
nuclei
velocity
umap_2
umap_1
G1
G2M
S
cell cycle
