## Additional file 2 for "Comparative framework and adaptation of ACME HS approach to single cell isolation from fresh-frozen endocrine tissues"

**Table S1: List of tissue samples (n=103) used in the study. Dissociation time, type of enzyme, cell viability and other dissociation parameters (RBCS, DRS) are indicated. Samples for which flow cytometry, RIN quantification and scRNA-seq was performed are marked.**

| sample number | method | type of tissue | enzyme | dissociation time (min) | RBCS (+-) | DRS (+-) | viability (%) | scRNA-seq | RNA extraction | flow cytometry |
|---|---|---|---|---|---|---|---|---|---|---|
| 1 | enzymatic | adrenal medullary tumor | MTDK | 20 | | + | 71 | + | | |
| 2 | enzymatic | adrenal medullary tumor | MTDK | 20 | + | + | 88 | + | | |
| 3 | enzymatic | adrenal medullary tumor | MTDK | 20 | + | + | 71 | + | | + |
| 4 | enzymatic | adrenal medullary tumor | MTDK | 20 | - | + | 89 | | | |
| 5 | enzymatic | adrenal medullary tumor | MTDK | 20 | - | + | 89 | | + | |
| 6 | enzymatic | adrenal medullary tumor | MTDK | 20 | - | + | 77 | | | |
| 7 | enzymatic | adrenal medullary tumor | MTDK | 20 | - | + | 50 | | | |
| 8 | enzymatic | adrenal medullary tumor | MTDK | 20 | - | + | 88 | | | |
| 9 | enzymatic | adrenal medullary tumor | MTDK | 20 | + | + | 95 | | | |
| 10 | enzymatic | adrenal medullary tumor | MTDK | 20 | + | + | 48 | | | |
| 11 | enzymatic | adrenal medullary tumor | Coll IV | 20 | + | + | 90 | | | |
| 12 | enzymatic | adrenal medullary tumor | MTDK | 25 | + | + | 69 | + | | |
| 13 | enzymatic | adrenal medullary tumor | MTDK | 25 | + | - | 78 | + | | |
| 14 | enzymatic | adrenal medullary tumor | MTDK | 25 | - | + | 61 | | | |
| 15 | enzymatic | adrenal medullary tumor | MTDK | 25 | + | - | 63 | | | |
| 16 | enzymatic | adrenal medullary tumor | MTDK | 25 | + | - | 65 | | | |
| 17 | enzymatic | adrenal medullary tumor | MTDK | 25 | + | + | 46 | | | |
| 18 | enzymatic | adrenal medullary tumor | NTDK | 25 | - | + | 74 | | | |
| 19 | enzymatic | adrenal medullary tumor | MTDK | 30 | + | - | 48 | | | |
| 20 | enzymatic | adrenal medullary tumor | MTDK | 30 | + | + | 35 | | | |
| 21 | enzymatic | adrenal medullary tumor | MTDK | 30 | + | + | 36 | | | |
| 22 | enzymatic | adrenal medullary tumor | MTDK | 30 | + | + | 59 | | | |
| 23 | enzymatic | adrenal medullary tumor | NTDK | 30 | + | - | 54 | | | |
| 24 | enzymatic | adrenal medullary tumor | NTDK | 30 | + | - | 33 | | | |
| 25 | enzymatic | adrenal medullary tumor | Coll IV | 30 | + | + | 58 | | | |
| 26 | enzymatic | adrenal medullary tumor | Coll IV | 30 | + | + | 53 | | | |
| 27 | ACME HS | adrenal medullary tumor | - | 60 | - | - | - | + | | |
| 28 | ACME HS | adrenal medullary tumor | - | 60 | - | - | - | + | | |
| 29 | ACME HS | adrenal medullary tumor | - | 60 | - | - | - | + | + | + |
| 30 | ACME HS | adrenal medullary tumor | - | 60 | - | - | - | + | | + |
| 31 | ACME HS | adrenal medullary tumor | - | 60 | - | - | - | + | | + |
| 32 | nuclei | adrenal medullary tumor | - | - | - | - | - | + | | |
| 33 | nuclei | adrenal medullary tumor | - | - | - | - | - | + | | |
| 34 | nuclei | adrenal medullary tumor | - | - | - | - | - | + | | |
| 35 | nuclei | adrenal medullary tumor | - | - | - | - | - | + | | |
| 36 | nuclei | adrenal medullary tumor | - | - | - | - | - | + | | |
| 37 | enzymatic | adrenocortical tumor | MTDK | 25 | | - | 77 | + | | |
| 38 | enzymatic | adrenocortical tumor | MTDK | 25 | + | - | 77 | + | | + |
| 39 | enzymatic | adrenocortical tumor | MTDK | 25 | + | + | 76 | + | | |
| 40 | enzymatic | adrenocortical tumor | Coll I | 25 | | + | 88 | + | | |

| | | | | | | | | | | |
|---|---|---|---|---|---|---|---|---|---|---|
| 41 | enzymatic | adrenocortical tumor | Coll I | 25 | | + | 87 | + | + | |
| 42 | enzymatic | adrenocortical tumor | MTDK | 30 | | - | 83 | | | |
| 43 | enzymatic | adrenocortical tumor | Coll IV | 30 | | + | 93 | | | |
| 44 | enzymatic | adrenocortical tumor | Coll IV | 30 | + | + | 72 | | | |
| 45 | enzymatic | adrenocortical tumor | Coll IV | 30 | + | + | 67 | | | |
| 46 | enzymatic | adrenocortical tumor | Coll I | 30 | | + | 77 | | | |
| 47 | ACME HS | adrenocortical tumor | - | 60 | - | - | - | + | | |
| 48 | ACME HS | adrenocortical tumor | - | 60 | - | - | - | + | + | + |
| 49 | ACME HS | adrenocortical tumor | - | 60 | - | - | - | + | | |
| 50 | ACME HS | adrenocortical tumor | - | 60 | - | - | - | + | | + |
| 51 | ACME HS | adrenocortical tumor | - | 60 | - | - | - | + | | + |
| 52 | nuclei | adrenocortical tumor | - | 60 | - | - | - | + | | |
| 53 | nuclei | adrenocortical tumor | - | 60 | - | - | - | + | | |
| 54 | enzymatic | thyroid carcinomas | MTDK | 20 | + | - | 56 | + | | |
| 55 | enzymatic | thyroid carcinomas | MTDK | 20 | | - | 87 | + | | |
| 56 | enzymatic | thyroid carcinomas | MTDK | 20 | | - | 71 | | | + |
| 57 | enzymatic | thyroid carcinomas | MTDK | 25 | | - | 64 | | + | |
| 58 | enzymatic | thyroid carcinomas | MTDK | 25 | + | - | 61 | | | |
| 59 | enzymatic | thyroid carcinomas | MTDK | 25 | + | - | 42 | + | | |
| 60 | enzymatic | thyroid carcinomas | MTDK | 30 | + | - | 69 | + | | |
| 61 | enzymatic | thyroid carcinomas | MTDK | 30 | | - | 51 | | | |
| 62 | enzymatic | thyroid carcinomas | MTDK | 30 | + | - | 75 | | | |
| 63 | ACME HS | thyroid carcinomas | - | 60 | - | - | - | | | + |
| 64 | ACME HS | thyroid carcinomas | - | 60 | - | - | - | | | + |
| 65 | ACME HS | thyroid carcinomas | - | 60 | - | - | - | + | + | + |
| 66 | enzymatic | PitNET | MTDK | 7 | | - | 91 | | | |
| 67 | enzymatic | PitNET | MTDK | 7 | | - | 61 | | | |
| 68 | enzymatic | PitNET | MTDK | 7 | | - | 61 | | | |
| 69 | enzymatic | PitNET | MTDK | 7 | | - | 59 | | | |
| 70 | enzymatic | PitNET | MTDK | 7 | + | - | 91 | + | | |
| 71 | enzymatic | PitNET | MTDK | 7 | + | + | 62 | | | |
| 72 | enzymatic | PitNET | MTDK | 7 | | - | 81 | | | + |
| 73 | enzymatic | PitNET | MTDK | 7 | | - | 39 | | + | |
| 74 | enzymatic | PitNET | Coll IV | 7 | + | - | 60 | | | |
| 75 | enzymatic | PitNET | Coll IV | 7 | + | - | 73 | | | |
| 76 | enzymatic | PitNET | MTDK | 7 | + | + | 71 | | | |
| 77 | enzymatic | PitNET | MTDK | 7 | - | + | 90 | | | |
| 78 | enzymatic | PitNET | MTDK | 7 | - | + | 98 | | | |
| 79 | enzymatic | PitNET | MTDK | 7 | - | - | 41 | | | |
| 80 | enzymatic | PitNET | Coll IV | 7 | - | + | 80 | | | |
| 81 | enzymatic | PitNET | Coll IV | 7 | - | - | 50 | + | | |
| 82 | enzymatic | PitNET | MTDK | 10 | - | - | 43 | | | |
| 83 | enzymatic | PitNET | MTDK | 10 | - | + | 94 | | | |
| 84 | enzymatic | PitNET | MTDK | 10 | - | + | 64 | | | |
| 85 | enzymatic | PitNET | MTDK | 10 | - | - | 63 | | | |
| 86 | enzymatic | PitNET | MTDK | 10 | - | - | 62 | | | |

| | | | | | | | | | | |
|---|---|---|---|---|---|---|---|---|---|---|
| 87 | enzymatic | PitNET | MTDK | 10 | - | - | 67 | | | |
| 88 | enzymatic | PitNET | MTDK | 10 | - | + | 44 | | | |
| 89 | enzymatic | PitNET | MTDK | 10 | - | - | 57 | | | |
| 90 | enzymatic | PitNET | MTDK | 10 | - | - | 87 | | | |
| 91 | enzymatic | PitNET | MTDK | 10 | - | - | 91 | | | |
| 92 | enzymatic | PitNET | MTDK | 10 | - | - | 77 | | | |
| 93 | enzymatic | PitNET | Coll IV | 10 | + | - | 84 | + | | |
| 94 | enzymatic | PitNET | MTDK | 15 | + | - | 60 | | | |
| 95 | enzymatic | PitNET | MTDK | 15 | - | - | 30 | | | |
| 96 | enzymatic | PitNET | MTDK | 15 | - | + | 45 | | | |
| 97 | enzymatic | PitNET | MTDK | 10 | - | - | 58 | + | | |
| 98 | enzymatic | PitNET | Coll IV | 10 | + | + | 65 | + | | |
| 99 | ACME HS | PitNET | - | 60 | - | - | - | | | + |
| 100 | ACME HS | PitNET | - | 60 | - | - | - | + | + | + |
| 101 | ACME HS | PitNET | - | 60 | - | - | - | + | | + |
| 102 | nuclei | PitNET | - | - | - | - | - | + | | |
| 103 | nuclei | PitNET | - | - | - | - | - | + | | |

***RBCS** - Red Blood Cell Lysis Solution (Miltenyi Biotec)
*__DRS__ - Dead Cell Removal Kit (Miltenyi Biotec)
*__MTDK__ - Multi Tissue Dissociation Kit (enzyme D) (Miltenyi Biotec)
*__NTDK__ - Neural Tissue Dissociation Kit P (enzyme A) (Miltenyi Biotec)
