## Additional file 3 for "Comparative framework and adaptation of ACME HS approach to single cell isolation from fresh-frozen endocrine tissues"

**Table S2: Enrichment score of key stress and death markers in ACME HS, enzyme and nuclei datasets. Genes associated with apoptosis, necrosis, cellular senescence, DNA damage, heat shock, unfolded protein response are represented. The percentage of all counts that belong to a given set of genes is indicated.**

| Signatures | Gene | ACME HS Adrenocortical tumors | ACME HS Adrenal medullary | ACME HS PitNETs | ACME HS Thyroid carcinomas | enzymatic Adrenocortical tumors | enzymatic Adrenal medullary | enzymatic PitNETs | enzymatic Thyroid carcinomas | nuclei Adrenocortical tumors | nuclei Adrenal medullary | nuclei PitNETs |
|---|---|---|---|---|---|---|---|---|---|---|---|---|
| necrosis | *HMGB1* | 0.04618426 | 0.06925339 | 0.04603636 | 0.04292872 | 0.06431831 | 0.05021457 | 0.05403836 | 0.03102966 | 0.01962648 | 0.0246764 | 0.02703135 |
| necrosis | *ATP5F1A* | 0.01639528 | 0.02120928 | 0.01711713 | 0.01370815 | 0.01932749 | 0.01204644 | 0.01852586 | 0.007779653 | 0.002485638 | 0.00376517 | 0.005442993 |
| necrosis | *CALR* | 0.01206507 | 0.01455296 | 0.02447213 | 0.01712569 | 0.01464967 | 0.01053026 | 0.01744666 | 0.005368599 | 0.001002307 | 0.0007080525 | 0.000895355 |
| necrosis | *ARHGAP45* | 0.0006516366 | 0.0002782426 | 0.0002534971 | 0.0001708521 | 0.001246524 | 0.00189937 | 0.003186213 | 0.0004272225 | 0.0004484307 | 0.0008543993 | 0.001044299 |
| necrosis | *S100A8* | 0.0008754062 | 0.0007002872 | 0.0008858887 | 0.0005424964 | 0.01021011 | 0.06009217 | 0.0002661078 | 0.0001912295 | 0.0001260035 | 13,2071900 | 0.0001039203 |
| necrosis | *S100A9* | 0.0009774266 | 0.0005222975 | 0.00046832 | 0.0005461348 | 0.01590388 | 0.0825104 | 0.0008668694 | 0.0002771594 | 0.0001145718 | 2,8891760 | 79,9743900 |
| necrosis | *NAMPT* | 0.005393698 | 0.003905071 | 0.008126661 | 0.002128693 | 0.008314448 | 0.01604655 | 0.00377837 | 0.004878602 | 0.01139982 | 0.005747684 | 0.001654621 |
| necrosis | *ANXA1* | 0.00316537 | 0.001924871 | 0.00110405 | 0.002208156 | 0.01298661 | 0.00667641 | 0.005159799 | 0.003749679 | 0.001294187 | 0.0007656829 | 0.0008960289 |
| necrosis | *KRT18* | 0.004131381 | 0.0006752998 | 0.01867394 | 0.02484178 | 0.001888852 | 60,4578900 | 0.007002793 | 0.005652412 | 0.000210997 | 46,1635800 | 0.0001673549 |
| necrosis | *TNF* | 6.16568800 | 0.0001123707 | 0.0002992218 | 24,7413400 | 0.0009514208 | 0.001247688 | 0.0005372212 | 0.000155815 | 8,4420360 | 0.0001283794 | 28,5488400 |
| necrosis | *AGER* | 0.0008315088 | 0.001081981 | 0.0004957528 | 0.001597231 | 0.0004360312 | 0.001030891 | 0.0006749694 | 0.0003703641 | 0.0003732168 | 0.0004553896 | 0.0003052366 |
| apoptosis | *CASP3* | 0.004102037 | 0.002668509 | 0.002389196 | 0.002380596 | 0.002454528 | 0.001967991 | 0.002843734 | 0.002213878 | 0.002316153 | 0.003444122 | 0.001548457 |
| apoptosis | *BAX* | 0.007686694 | 0.00145592 | 0.002413642 | 0.0007907977 | 0.009760695 | 0.006003619 | 0.008829019 | 0.001846491 | 0.001514232 | 0.001030531 | 0.001102914 |
| apoptosis | *BAD* | 0.007488401 | 0.004215078 | 0.004263085 | 0.00252189 | 0.005488453 | 0.002202771 | 0.00331075 | 0.00182314 | 0.0003633803 | 0.00029761 | 0.0005376017 |
| apoptosis | *BID* | 0.0006377724 | 0.0004145278 | 0.0004617181 | 0.000471882 | 0.003702875 | 0.00273005 | 0.0006874142 | 0.002045889 | 0.000604245 | 0.00090257 | 0.0004847641 |
| apoptosis | *APAF1* | 0.001872543 | 0.001932321 | 0.0008598655 | 0.00238042 | 0.002020049 | 0.002868489 | 0.002216316 | 0.00251553 | 0.003540457 | 0.002086184 | 0.002082814 |
| apoptosis | *TP53* | 0.001601209 | 0.001400935 | 0.0009738565 | 0.001353208 | 0.001106504 | 0.001509634 | 0.001939501 | 0.001114386 | 0.001387923 | 0.001790445 | 0.0008629755 |
| apoptosis | *FAS* | 0.00288045 | 0.0008602322 | 46,3735700 | 0.0002488027 | 0.002305044 | 0.000720613 | 0.000670497 | 0.0004185902 | 0.003712661 | 0.001018499 | 0.0005644215 |
| apoptosis | *TNFRSF10B* | 0.002974873 | 0.0004840692 | 0.0001982579 | 0.0006137563 | 0.001854898 | 0.0009699938 | 0.0008965998 | 0.001879143 | 0.004000981 | 0.001198636 | 0.0008131107 |
| apoptosis | *CYCS* | 0.01096457 | 0.01326353 | 0.007512603 | 0.003950346 | 0.01246605 | 0.008718624 | 0.008323618 | 0.001957239 | 0.0004215813 | 0.0009231766 | 0.0004580033 |
| apoptosis | *BCL2* | 0.00025316 | 0.0025149 | 0.0009362198 | 0.02705494 | 0.001226855 | 0.004296135 | 0.005153167 | 0.08561267 | 0.003879408 | 0.02046997 | 0.01531816 |
| apoptosis | *AIFM1* | 0.003171797 | 0.002187386 | 0.001094593 | 0.001606648 | 0.003312552 | 0.001016385 | 0.002013475 | 0.001528855 | 0.004308583 | 0.002309694 | 0.001644949 |
| DNA_dmg | *TP53* | 0.001601209 | 0.001400935 | 0.0009738565 | 0.001353208 | 0.001106504 | 0.001509634 | 0.001939501 | 0.001114386 | 0.001387923 | 0.001790445 | 0.0008629755 |
| DNA_dmg | *BRCA1* | 0.001810897 | 0.00151125 | 0.002017889 | 0.002208233 | 0.0006925573 | 0.002170925 | 0.00129066 | 0.001164589 | 0.0009858727 | 0.001565121 | 0.002153529 |
| DNA_dmg | *CHEK2* | 0.001419915 | 0.0001694754 | 0.0002387893 | 0.000275431 | 0.001037202 | 0.00027672 | 0.0002566969 | 0.0003669098 | 0.001398243 | 0.0005486549 | 0.0006775394 |
| DNA_dmg | *ATM* | 0.01116395 | 0.02656568 | 0.02112073 | 0.02806129 | 0.008123506 | 0.02673727 | 0.02968061 | 0.01765452 | 0.01893455 | 0.02632923 | 0.02843498 |
| DNA_dmg | *RAD51* | 0.0003536354 | 0.0001309483 | 0.0001131968 | 67,3080400 | 0.0001860613 | 0.0001092717 | 8.899711e-05 | 69,0511700 | 0.0001172027 | 0.0001235596 | 0.0001118159 |
| DNA_dmg | *RPA1* | 0.002071312 | 0.002184553 | 0.003931805 | 0.001787569 | 0.001377315 | 0.001549193 | 0.003360275 | 0.003004048 | 0.003002165 | 0.003512794 | 0.005925825 |
| DNA_dmg | *MDM2* | 0.01559148 | 0.00636195 | 0.003003378 | 0.006050141 | 0.007405273 | 0.01034456 | 0.009611315 | 0.005530033 | 0.01212116 | 0.006307162 | 0.006021719 |
| DNA_dmg | *ATR* | 0.02132393 | 0.01022471 | 0.007885424 | 0.01396389 | 0.01101679 | 0.01348956 | 0.008828519 | 0.01251285 | 0.03779912 | 0.008179073 | 0.01763599 |
| DNA_dmg | *XRCC5* | 0.009670555 | 0.02183485 | 0.0113063 | 0.01580023 | 0.01599494 | 0.01235315 | 0.0126527 | 0.01766393 | 0.01493604 | 0.01916674 | 0.01440894 |
| unfolded | *ATF4* | 0.01313637 | 0.01345351 | 0.007321867 | 0.00494761 | 0.00738441 | 0.009408122 | 0.009027123 | 0.002809338 | 0.004281262 | 0.002459752 | 0.002724193 |
| unfolded | *ATF6* | 0.00356311 | 0.006129374 | 0.004792936 | 0.01214166 | 0.003227391 | 0.004483093 | 0.005016875 | 0.01599231 | 0.01653325 | 0.01261998 | 0.01341841 |
| unfolded | *XBP1* | 0.003266255 | 0.005420776 | 0.005004801 | 0.01875834 | 0.003543933 | 0.004214783 | 0.006414593 | 0.006535581 | 0.00196404 | 0.001464567 | 0.002214107 |
| unfolded | *HSPA5* | 0.01488166 | 0.03170209 | 0.03094525 | 0.03354565 | 0.01935127 | 0.01630661 | 0.03585573 | 0.01204541 | 0.0008032513 | 0.000854616 | 0.001534223 |
| unfolded | *DDIT3* | 0.03068386 | 0.01022668 | 0.00436529 | 0.001516282 | 0.0100865 | 0.004059849 | 0.004872548 | 0.0008042045 | 0.001802302 | 0.001037384 | 0.0004011718 |
| unfolded | *HERPUD1* | 0.01350852 | 0.01267367 | 0.007485925 | 0.007730957 | 0.02589918 | 0.01282864 | 0.01088636 | 0.006625201 | 0.005139913 | 0.004078085 | 0.001836082 |
| unfolded | *DNAJC3* | 0.005641139 | 0.01655151 | 0.00476497 | 0.01659218 | 0.006679985 | 0.006425159 | 0.0126971 | 0.01135887 | 0.008675875 | 0.007816119 | 0.007452788 |

| | | | | | | | | | | | | |
|---|---|---|---|---|---|---|---|---|---|---|---|---|
| unfolded | *ERN1* | 0.008869876 | 0.00169557 | 0.001271614 | 0.005375525 | 0.007697833 | 0.001633505 | 0.004911937 | 0.006442162 | 0.008614007 | 0.002734061 | 0.005329857 |
| unfolded | *ERN2* | 0.0001533555 | 22,7179700 | 37,9822100 | 9,8779610 | 14,8555700 | 0,2513300 | 0.0001001301 | 0,7986350 | 0,0139100 | 16,2634800 | 38,1903900 |
| unfolded | *PDIA6* | 0.01148829 | 0.007014642 | 0.009712996 | 0.01231216 | 0.02385102 | 0.005892162 | 0.010329 | 0.008692163 | 0.002241486 | 0.001548908 | 0.001802533 |
| senescence | *CDKN1A* | 0.009883957 | 0.002818875 | 0.001215011 | 0.004259349 | 0.002781881 | 0.003487497 | 0.004350714 | 0.0007546974 | 0.0003566668 | 0.000761582 | 0.0003017513 |
| senescence | *CDKN2A* | 0.0007219538 | 0.001255448 | 0.005104228 | 0.0001550224 | 0.009722765 | 0.0005134505 | 0.004739099 | 0.000427346 | 0.001006728 | 0.000545364 | 0.004591217 |
| senescence | *IGFBP3* | 0.01143321 | 0.001754264 | 0.0006319229 | 0.0007010089 | 0.003325028 | 0.001899722 | 0.0006448783 | 0.0004035439 | 0.0002470814 | 0.0005654194 | 0.0003312915 |
| senescence | *GADD45A* | 0.004647787 | 0.001284588 | 0.002579488 | 0.00122829 | 0.001983614 | 0.001660693 | 0.002840425 | 0.0007116607 | 0.0001699023 | 0.0003784129 | 0.0003012702 |
| senescence | *CCND1* | 0.005626422 | 0.00810493 | 0.005958748 | 0.02003892 | 0.004581528 | 0.008438807 | 0.009665745 | 0.00832921 | 0.002004818 | 0.001620206 | 0.001842544 |
| senescence | *CDKN2B* | 0.0001954046 | 0.0005153939 | 0.0005777793 | 0.0001189456 | 0.0004864411 | 0.0003072455 | 0.001384868 | 92,5469200 | 0.0001417967 | 95,3742900 | 0.0002060789 |
| senescence | *IL1A* | 34,57272000 | 15,3971700 | 7,4904720 | 0,0000000 | 35,0349500 | 19,2792900 | 1.566157e-05 | 0.0001031494 | 6,1815930 | 2,6121660 | 22,3454100 |
| senescence | *IL1B* | 35,60605000 | 85,9113100 | 0.0001783432 | 0,0000000 | 0.001581581 | 0.001777343 | 0.0002365738 | 0.0003897463 | 19,9720500 | 0.0001139469 | 0.0002109411 |
| senescence | *IL6* | 4,24886300 | 30,6130500 | 62,9798600 | 13,4692700 | 44,8035900 | 56,9400800 | 1.258767e-05 | 37,5768400 | 1,5150500 | 81,5127300 | 10,3249300 |
| senescence | *IL10* | 11,70352000 | 95,3677200 | 0.0003941419 | 0,0000000 | 0.0001945992 | 0.000329775 | 7.243003e-05 | 38,5131400 | 9,5265630 | 38,1874800 | 6,0668390 |
| senescence | *HMGA1* | 0.0003070307 | 0.001866823 | 0.0009392061 | 0.001495973 | 0.0009357762 | 0.001925468 | 0.002824355 | 0.0005623773 | 62,5948400 | 0.0003742615 | 0.0001372323 |
| senescence | *HMGB2* | 0.007633135 | 0.00591744 | 0.004780525 | 0.001471903 | 0.009739406 | 0.007077975 | 0.005553095 | 0.0009865863 | 0.0003858 | 0.0006518024 | 0.0008140395 |
| senescence | *UBB* | 0,09020205 | 0.05711255 | 0.05220257 | 0.03416852 | 0.08357375 | 0.02588403 | 0.04328713 | 0.02794551 | 0.0001095723 | 44,2903700 | 0.0005937678 |
| heat_shock | *HSP90AA1* | 0,05864521 | 0,0853695 | 0,0283317 | 0,0872833 | 0,1101236 | 0,0679252 | 4.64037e-06 | 0,0382150 | 0,0116579 | 0,0124983 | 0,0091123 |
| heat_shock | *HSP90AB1* | 0,06434839 | 0,1281465 | 0,0401991 | 0,0684396 | 0,1864129 | 0,0711521 | 0.000182865 | 0,0326688 | 0,0081196 | 0,0084698 | 0,0058650 |
| heat_shock | *HSP90B1* | 0,04006888 | 0,0513432 | 0,0307673 | 0,0996798 | 0,0506602 | 0,0281267 | 0.004179348 | 0,0222492 | 0,0095600 | 0,0045288 | 0,0036748 |
| heat_shock | *HSPA12A* | 0,01036174 | 0,0047684 | 0,0085436 | 0,0020504 | 0,0020034 | 0,0057722 | 0.0006228466 | 0,0061601 | 0,0180535 | 0,0105358 | 0,0103876 |
| heat_shock | *HSPA12B* | 0,00005809 | 0,0000647 | 0,0000530 | 0,0000638 | 0,0000318 | 0,0001525 | 0.01435398 | 0,0001057 | 0,0001878 | 0,0001365 | 0,0000199 |
| heat_shock | *HSPA13* | 0,00359016 | 0,0084026 | 0,0049343 | 0,0100750 | 0,0022336 | 0,0033255 | 0.00955298 | 0,0015540 | 0,0005091 | 0,0010009 | 0,0010543 |
| heat_shock | *HSPA14* | 0,00138321 | 0,0008407 | 0,0006187 | 0,0008893 | 0,0004276 | 0,0011491 | 0.001835134 | 0,0002971 | 0,0002806 | 0,0003397 | 0,0003192 |
| heat_shock | *HSPA14.1* | 0,00389819 | 0,0024863 | 0,0014930 | 0,0013341 | 0,0010515 | 0,0016079 | 0.008631792 | 0,0010241 | 0,0013227 | 0,0011629 | 0,0009006 |
| heat_shock | *HSPA1A* | 0,01369285 | 0,0058262 | 0,0017086 | 0,0021062 | 0,0152034 | 0,0218774 | 8.033676e-05 | 0,0080272 | 0,0037226 | 0,0004472 | 0,0000283 |
| heat_shock | *HSPA1B* | 0,00406424 | 0,0034534 | 0,0007824 | 0,0005366 | 0,0029149 | 0,0103168 | 0.006750223 | 0,0046912 | 0,0026665 | 0,0004026 | 0,0002561 |
| heat_shock | *HSPA1L* | 0,00010311 | 0,0001411 | 0,0001387 | 0,0001293 | 0,0000685 | 0,0001932 | 0.0117732 | 0,0000793 | 0,0000737 | 0,0001573 | 0,0001072 |
| heat_shock | *HSPA2* | 0,00043316 | 0,0003966 | 0,0000922 | 0,0006063 | 0,0002658 | 0,0001553 | 0.0004282073 | 0,0002833 | 0,0000803 | 0,0001328 | 0,0000935 |
| heat_shock | *HSPA4* | 0,00677681 | 0,0106934 | 0,0050304 | 0,0060583 | 0,0067349 | 0,0044088 | 0.01058081 | 0,0070779 | 0,0070654 | 0,0060508 | 0,0054295 |
| heat_shock | *HSPA4L* | 0,00820974 | 0,0074314 | 0,0081686 | 0,0029460 | 0,0025498 | 0,0075774 | 0.009596654 | 0,0042788 | 0,0055526 | 0,0080850 | 0,0058941 |
| heat_shock | *HSPA5* | 0,01624317 | 0,0317338 | 0,0209975 | 0,0335457 | 0,0195496 | 0,0168597 | 0.109897 | 0,0110114 | 0,0007994 | 0,0009698 | 0,0015859 |
| heat_shock | *HSPA6* | 0,00012688 | 0,0007364 | 0,0000682 | 0,0000791 | 0,0009267 | 0,0022013 | 0.01455451 | 0,0004664 | 0,0000994 | 0,0009335 | 0,0000744 |
| heat_shock | *HSPA8* | 0,02244468 | 0,0286931 | 0,0155037 | 0,0207882 | 0,0322779 | 0,0271291 | 0.03585573 | 0,0082239 | 0,0010577 | 0,0012422 | 0,0008485 |
| heat_shock | *HSPA9* | 0,02492587 | 0,0207465 | 0,0093126 | 0,0095092 | 0,0192189 | 0,0149980 | 0.0008835194 | 0,0068835 | 0,0171281 | 0,0088267 | 0,0037809 |
| heat_shock | *HSPB1* | 0,05537426 | 0,0203033 | 0,0035135 | 0,0207960 | 0,1061258 | 0,0170496 | 0.001356661 | 0,0112772 | 0,0040265 | 0,0005853 | 0,0001305 |
| heat_shock | *HSPB11* | 0,00814652 | 0,0045818 | 0,0046523 | 0,0023746 | 0,0088743 | 0,0029966 | 0.003852043 | 0,0045194 | 0,0043755 | 0,0028586 | 0,0036362 |
| heat_shock | *HSPB2* | 0,00008531 | 0,0002130 | 0,0000113 | 0,0000960 | 0,0000438 | 0,0000928 | 1.32987e-05 | 0,0000749 | 0,0000549 | 0,0000353 | 0,0000070 |
| heat_shock | *HSPB3* | 0,00000086 | 0,0000237 | 0,0035792 | 0,0000000 | 0,0000000 | 0,0000016 | 0.03392874 | 0,0000097 | 0,0000000 | 0,0000004 | 0,0000000 |
| heat_shock | *HSPB6* | 0,00943175 | 0,0019739 | 0,0000345 | 0,0005364 | 0,0078670 | 0,0046095 | 0.03780706 | 0,0000588 | 0,0005351 | 0,0001796 | 0,0000180 |
| heat_shock | *HSPB7* | 0,00199766 | 0,0002503 | 0,0000535 | 0,0000974 | 0,0017329 | 0,0015409 | 0.000688174 | 0,0001736 | 0,0004208 | 0,0008486 | 0,0000048 |
| heat_shock | *HSPB8* | 0,00290347 | 0,0010430 | 0,0002334 | 0,0008296 | 0,0010276 | 0,0004921 | 0.01545384 | 0,0023739 | 0,0004821 | 0,0003749 | 0,0002058 |
| heat_shock | *HSPB9* | 0,00000724 | 0,0000207 | 0,0000000 | 0,0003233 | 0,0000029 | 0,0000045 | 0.0005614918 | 0,0000469 | 0,0000057 | 0,0000169 | 0,0000000 |
| heat_shock | *HSPBAP1* | 0,00120608 | 0,0010166 | 0,0022722 | 0,0013691 | 0,0012693 | 0,0023479 | 0.08973372 | 0,0025779 | 0,0046224 | 0,0026915 | 0,0051700 |
| heat_shock | *HSPBP1* | 0,00205699 | 0,0018138 | 0,0013459 | 0,0021569 | 0,0016311 | 0,0008950 | 6.01026e-06 | 0,0009463 | 0,0006114 | 0,0008544 | 0,0009444 |

| | | | | | | | | | | | | |
|---|---|---|---|---|---|---|---|---|---|---|---|---|
| heat_shock | *HSPD1* | 0,12540340 | 0,0275582 | 0,0118026 | 0,0192617 | 0,1203400 | 0,0176143 | 0.0002708268 | 0,0117450 | 0,0150033 | 0,0056482 | 0,0029607 |
| heat_shock | *HSPE1* | 0,27823903 | 0,0362301 | 0,0184628 | 0,0130209 | 0,1323436 | 0,0128363 | 0.001913361 | 0,0053618 | 0,0033325 | 0,0007067 | 0,0002988 |
| heat_shock | *HSPG2* | 0,00930337 | 0,0020348 | 0,0009466 | 0,0018360 | 0,0085369 | 0,0040430 | 1.271002e-05 | 0,0023849 | 0,0158057 | 0,0029399 | 0,0003105 |
| heat_shock | *HSPH1* | 0,00839267 | 0,0156257 | 0,0100425 | 0,0216913 | 0,0058957 | 0,0146988 | 0.006944365 | 0,0210549 | 0,0058588 | 0,0079179 | 0,0041470 |
